# T-ALPHA: A Hierarchical Transformer-Based Deep Neural Network for Protein-Ligand Binding Affinity Prediction With Uncertainty-Aware Self-Learning for Protein-Specific Alignment

**DOI:** 10.1101/2024.12.19.629497

**Authors:** Gregory W. Kyro, Anthony M. Smaldone, Yu Shee, Chuzhi Xu, Victor S. Batista

**Affiliations:** Department of Chemistry, Yale University

## Abstract

There is significant interest in targeting disease-causing proteins with small molecule inhibitors to restore healthy cellular states. The ability to accurately predict the binding affinity of small molecules to a protein target in silico enables the rapid identification of candidate inhibitors and facilitates the optimization of on-target potency. In this work, we present T-ALPHA, a novel deep learning model that enhances protein-ligand binding affinity prediction by integrating multimodal feature representations within a hierarchical transformer framework to capture information critical to accurately predicting binding affinity. T-ALPHA outperforms all existing models reported in the literature on multiple benchmarks designed to evaluate protein-ligand binding affinity scoring functions. Remarkably, T-ALPHA maintains state-of-the-art performance when utilizing predicted structures rather than crystal structures, a powerful capability in real-world drug discovery applications where experimentally determined structures are often unavailable or incomplete. Additionally, we present an uncertainty-aware self-learning method for protein-specific alignment that does not require additional experimental data, and demonstrate that it improves T-ALPHA’s ability to rank compounds by binding affinity to biologically significant targets such as the SARS-CoV-2 main protease and the epidermal growth factor receptor. To facilitate implementation of T-ALPHA and reproducibility of all results presented in this paper, we have made all of our software available at https://github.com/gregory-kyro/T-ALPHA.

## 1 Introduction

There is growing scientific and societal interest in developing therapeutic interventions targeting diseases that disproportionately affect humans and remain inadequately addressed by current medical treatments.^1^ Protein dysregulation, including overexpression and aberrant post-translational modifications, is a fundamental factor in the pathogenesis of numerous human diseases. For instance, in neurodegenerative disorders like Alzheimer’s and Parkinson’s diseases, abnormal protein modifications can lead to aggregation and neuronal death.^2^ Similarly, in cancer, the overexpression of certain proteins disrupts cellular homeostasis, contributing to uncontrolled cell proliferation.^3^ Targeting these dysregulated proteins with small molecules (i.e., ligands) offers a promising therapeutic strategy to restore normal cellular function and transition cells from a disease state to a healthy state.^4^

A typical pipeline to develop therapeutic compounds consists of several sequential stages: target identification (selecting a biological target linked to disease),^5^ target validation (confirming the target’s role in disease progression and suitability for therapeutic intervention),^6^ hit identification (screening for compounds with initial activity against the target),^7^ lead optimization (refining compounds for potency, selectivity, and desirable pharmacokinetics),^8^ preclinical testing (evaluating safety and efficacy in non-human models),^9^ and clinical development (testing for safety and efficacy in human trials).^10^ In this work, we are focused on the hit identification and lead optimization stages, specifically, predicting protein-ligand binding affinity.

Machine learning (ML) has been widely applied to protein-ligand binding affinity prediction and has become central to computer aided-drug design more broadly, with applications including protein structure prediction,^11, 12^ molecular docking,^13^ small molecule property prediction,^14, 15^ and others.^16^ Traditional ML approaches for protein-ligand binding affinity prediction, such as random forests^17–20^ and shallow neural networks,^21^ are increasingly being replaced by deep learning methods that are better suited for learning geometric representations of molecular structures.^22^ Examples include convolutional neural networks (CNNs) which utilize convolutional operations to capture local spatial features from voxelized grids,^23–35^ graph neural networks (GNNs) which employ message passing to model relational information in molecular graphs,^33, 35–58^ and, more recently, transformers which utilize self- and cross-attention to model long-range dependencies within and between embeddings, respectively.^59–82^ Moreover, it has been demonstrated that transformer-based multimodal feature representation learning of proteins is effective for extracting features from the protein that are important for predicting protein-ligand binding affinity,^73^ a finding that has inspired multiple components of our work.

We present T-ALPHA, a novel deep learning model for protein-ligand binding affinity prediction, designed to comprehensively integrate multimodal data representations and leverage hierarchical transformer mechanisms to capture the intricate physicochemical, structural, and spatial dynamics governing protein-ligand binding interactions. T-ALPHA processes input data through three distinct channels, corresponding to the protein, ligand, and protein-ligand complex. Each channel independently learns a rich and optimized feature representation for its specific component, ensuring that critical properties of proteins, ligands, and their interactions are captured.

Within the protein and ligand channels, cross-attention mechanisms are employed to integrate complementary information between diverse feature representations. The protein channel combines features derived from the protein’s surface topography and curvature using point cloud-based quasi-geodesic convolutions, connectivity-based structural information from an E(*n*) equivariant graph neural network (EGNN), and sequence-derived evolutionary and structural embeddings derived from a pretrained transformer-based model. Similarly, the ligand channel integrates molecular-level physicochemical descriptors, graph-based structural features from an E(*n*) EGNN, and relational information extracted from SMILES strings using a pretrained transformer encoder. The protein-ligand complex channel focuses exclusively on modeling the interactions between the protein and ligand through an E(*n*) EGNN, capturing spatial relationships and interaction-specific features essential for binding affinity prediction. After processing through these channels, an additional layer of cross-attention integrates the outputs from the protein, ligand, and protein-ligand complex channels, enabling the model to combine their complementary perspectives into a unified, hierarchical representation. This design ensures that T-ALPHA effectively models the complex dependencies between proteins and ligands, resulting in state-of-the-art performance across multiple benchmarks.

Typically, deep learning models for protein-ligand binding affinity prediction are trained and benchmarked on datasets containing many different proteins to assess generalization across diverse targets. While this is valuable, in most practical applications, the focus is on a single, disease-relevant protein, where high accuracy for that specific target is paramount. Current methods for target-specific alignment, such as active learning-based approaches,^83^ require acquisition of additional experimental data which can be resource-intensive and slow. In this work, we introduce a novel uncertainty-aware self-learning method that enables protein-specific alignment without the need for new experimental data. Applied to two distinct protein targets, namely SARS-CoV-2 main protease (Mpro) and the epidermal growth factor receptor (EGFR), our approach improves the model’s target-specific ranking of compounds by binding affinity, offering a resource-efficient strategy for real-world applications that require high accuracy on defined protein targets.

The ability of T-ALPHA to effectively model protein-ligand interactions, combined with the proposed self-learning method for protein-specific alignment, underscores the utility of this work to both broad and target-focused applications in protein-ligand binding affinity prediction.

## 2 Data

### 2.1 Data Curation

#### 2.1.1 Protein-Ligand Binding Affinity Data

T-ALPHA is trained on experimentally determined protein-ligand complex structures deposited in the Protein Data Bank, each labeled with a binding affinity measurement. The PDBbind database,^84–86^ widely regarded as the primary data source for evaluating deep learning models for protein-ligand binding affinity prediction, contains 19_443 protein-ligand complex structures in its latest update (v2020), the majority of which are X-ray crystal structures, with a smaller portion determined via Nuclear Magnetic Resonance (NMR) spectroscopy. Each binding affinity value is reported as either an inhibition constant (*K*_i_), dissociation constant (*K*_d_), or half-maximal inhibitory concentration (IC_50_).

Because the biochemical assays performed to obtain these measurements operate within specific dynamic ranges, some measurements are reported with inequalities (“>”, “<”) rather than exact (“=”) or approximate (“∼”) values. While most protein-ligand binding affinity prediction work treats all labels as exact values (i.e., assuming the operator to be “=” for each datapoint), we consider the original operators via a custom loss function that we constructed to more accurately penalize model predictions during training and validation (for more details, see Section 4.1).

The complete v2020 release of the PDBbind database, referred to as the *general set*, is filtered into a *refined set* (5_316 entries),^87^ which comprises relatively high-quality crystal structures with relatively reliable binding affinity values. The refined set is further filtered into the *core set* (290 entries), representing the highest quality datapoints, with 58 proteins, each co-crystallized with five different ligands. Moreover, each binding affinity measurement is reported as an exact value (“=”). The core set serves as the basis for the Comparative Assessment of Scoring Functions (CASF) 2016 benchmark,^88^ which is the most widely used benchmark for assessing protein-ligand binding affinity scoring functions.

While the CASF 2016 benchmark is widely regarded as the gold standard of the field, it is unable to adequately assess the ability of models to generalize to protein-ligand complexes with low similarity to those found in the rest of the PDBbind database (i.e., the corresponding training and validation data).^89^ Specifically, the majority of the datapoints in the core set contain either identical proteins or chemically similar ligands with entries in the rest of the PDBbind v2020 general set, introducing data leakage that inflates performance estimates on the CASF 2016 benchmark with respect to generalizability to low-similarity protein-ligand complexes. In this work, we utilize the CASF 2016 test set exclusively to compare the T-ALPHA architecture to the many models reported in the literature that also benchmark on this dataset.

To address some of the limitations of the CASF 2016 benchmark for assessing protein-ligand binding affinity scoring functions, Leak Proof PDBbind (LP-PDBbind) was developed to minimize protein sequence and ligand structure similarities across training, validation, and test sets, while also ensuring distinct protein-ligand structural interaction patterns across these sets.^90^ Ligand similarity was calculated as the Dice similarity^91^ between 1024-bit Morgan fingerprints, and protein similarity was calculated as the sequence identity between Needleman-Wunsch-aligned^92^ amino acid sequences. The splitting approach utilized ensures that no training datapoints have a protein similarity or ligand similarity greater than 0.5 or 0.99, respectively, to any datapoint in the validation or test sets, and that no validation datapoints have a protein similarity or ligand similarity greater than 0.9 or 0.99, respectively, to any datapoint in the test set. Additionally, proteo-chemometric interaction fingerprints^93^ were used to evaluate interaction patterns. These fingerprints extend the Morgan fingerprint by incorporating the spatial interactions between ligand atoms and adjacent protein residues, mapping interaction patterns to a fixed-size integer vector with a length of 256. Pairwise interaction fingerprint similarities were then calculated using a weighted Tanimoto similarity score.^94^ The data splitting protocol ensures distinct separation of training, validation and test data in terms of these interaction fingerprints.

In addition to the LP-PDBbind split, the same research group developed a test set for benchmarking protein-ligand binding affinity scoring functions that is independent of the data contained in the PDBbind database, referred to as the BDB2020+ dataset. This dataset consists of datapoints from the BindingDB database^95^ that were deposited after 2020, reducing the potential for data leakage with v2020 of the PDBbind. While BindingDB provides experimental binding affinity data for protein-ligand complexes, it does not include the corresponding experimentally determined structures. To address this, binding affinity data from BindingDB was cross-referenced with matching experimental structures in the RCSB Protein Data Bank.^96^

Additionally, two protein-specific test sets were developed by the same research group and employed in this work that focus on the SARS-CoV-2 main protease (Mpro)^97^ and the epidermal growth factor receptor (EGFR),^98^ a receptor tyrosine kinase implicated in multiple types of cancer. The Mpro dataset contains 40 data points, each with a unique ligand co-crystallized with Mpro, while the EGFR dataset comprises 23 data points, each with a unique ligand co-crystallized with EGFR; all data points are labeled with experimentally determined binding affinity measurements. For full details on the preparation and validation of the LP-PDBbind splits, the BDB2020+ test set, as well as the Mpro and EGFR test sets, please refer to the original paper.^90^

#### 2.1.2 Ligand SMILES Transformer Pretraining Data

In T-ALPHA, one of the ligand representations is a feature vector that is extracted from a transformer encoder pretrained for masked token prediction on a dataset of SMILES strings. For pretraining, we utilize a comprehensive dataset that we have previously curated,^99^ which combines all of the SMILES strings from ChEMBL 33 (∼ 2.4 million bioactive molecules with drug-like properties),^100^ GuacaMol v1 (∼ 1.6 million molecules derived from ChEMBL 24 that have been synthesized and tested against biological targets),^101^ MOSES (∼ 1.8 million molecules selected from ZINC 15 to maximize internal diversity and suitability for medicinal chemistry),^102^ BindingDB (∼ 1.2 million unique small molecules bound to proteins),^95^ and the v2020 release of the PDBbind general set (15_710 unique small molecules bound to proteins).^84–86^

#### 2.1.3 Predicted Protein-Ligand Complex Structures

For each of the test sets employed in this work, we use Chai1, a state-of-the-art multimodal foundation model for molecular structure prediction, to predict the protein-ligand complex 3D structure from each protein amino acid sequence and ligand SMILES string pair. In the context of this work, we utilize predicted structures to assess the robustness of T-ALPHA in scenarios where experimentally determined structures are unavailable or incomplete. We provide full implementation details of Chai1 in Section 9.

### 2.2 Data Preparation

#### 2.2.1 Protein-Ligand Complex Structures

We preprocessed all protein-ligand complex structures, including both the experimentally determined and predicted structures. Solvent molecules were removed using Biopython^103^ to allow the model to implicitly learn solvent effects. Each protein was corrected for missing heavy atoms and residues using PDBFixer and OpenMM.^104^ We then added missing hydrogen atoms to the proteins and ligands with Open Babel.^105^ For each datapoint, we extracted the protein pocket by taking any residue in the protein that contains at least one atom that is within 8 Å of at least one ligand non-hydrogen atom.

#### 2.2.2 Ligand SMILES Transformer Pretraining Data

To prepare the pretraining dataset for the SMILES-based transformer encoder that we use as a ligand feature extractor, we processed each SMILES string using RDKit^106^ by creating a mol object and canonicalizing. After removing duplicate canonical SMILES and those that failed processing, 5_131_118 valid and unique SMILES strings remained out of the original 5_791_565 entries. Tokenization resulted in a vocabulary of 379 unique tokens, of which only 132 tokens occur in the PDBbind v2020 general set. We removed any SMILES string that contains at least one token that does not occur in any datapoint in the PDBbind v2020 general set, resulting in 4_810_575 remaining SMILES strings and a significant reduction of the vocabulary. In order to significantly increase computational efficiency, we removed any SMILES string with more than the 95^th^ percentile number of tokens, reducing the block size from 385 to 155 tokens. This entire preparation yielded a pretraining dataset of 4_778_512 datapoints.

### 2.3 Data Featurization

#### 2.3.1 Protein Featurization

In T-ALPHA, there are three channels of data processing, corresponding to the protein, ligand, and protein-ligand complex. For the protein channel, we represent and process the protein in three distinct ways: (1) a sparse graph derived from the protein pocket which is processed by an E(*n*) EGNN to learn connectivity-based structural features, (2) a point cloud of the surface of the protein pocket which is processed via quasi-geodesic convolutions to learn features pertaining to the surface topography and curvature, and (3) an amino acid sequence-derived embedding which is processed by a multilayer perceptron (MLP) to learn complementary global evolutionary and structural information (Figure 1).

**Figure 1.**
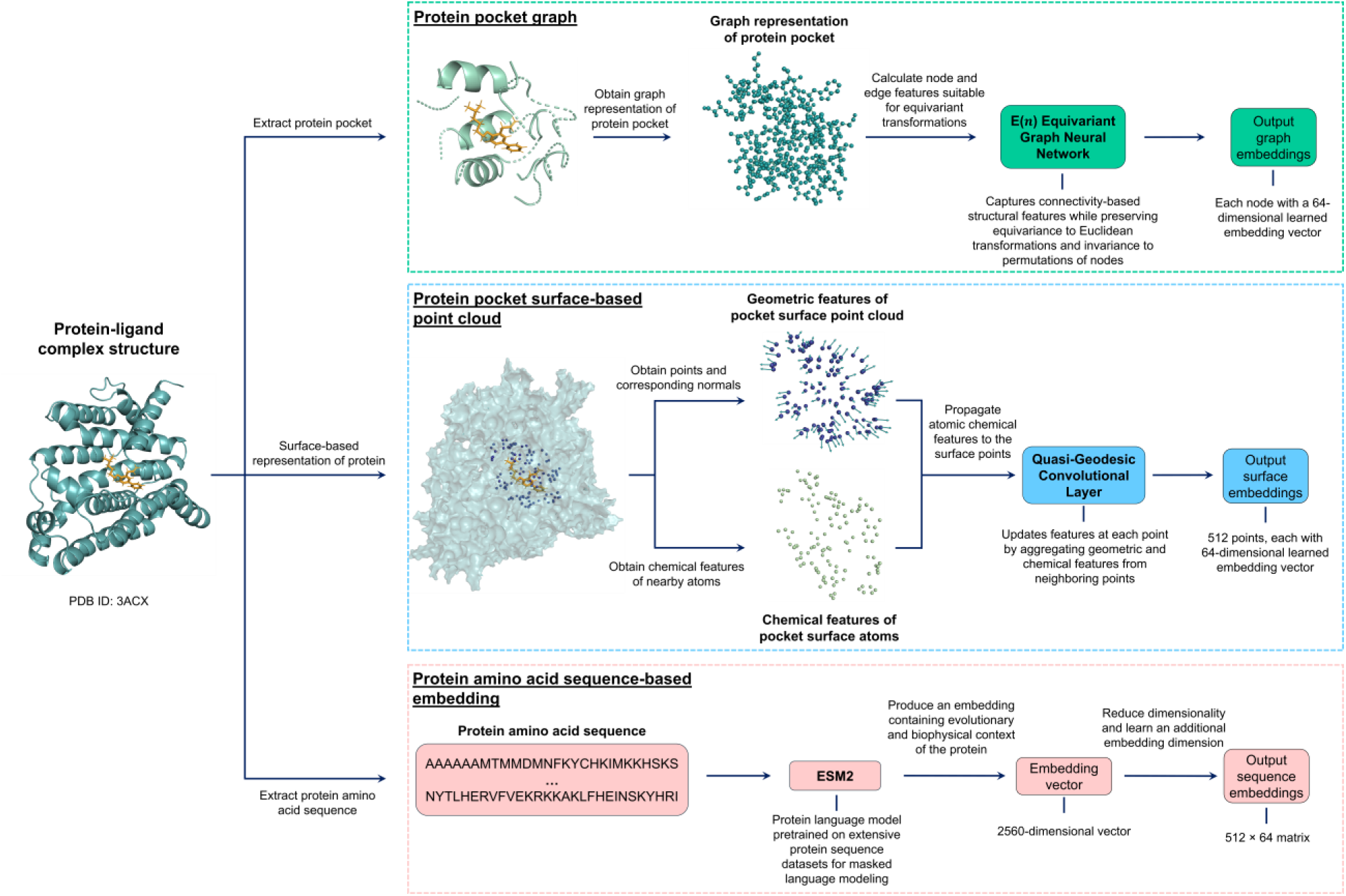
The protein channel of the T-ALPHA pipeline. The protein is represented and processed in three distinct ways: (1) a graph representation of the protein pocket (top, green box), where nodes and edges are annotated with atomic and chemical features, is processed by an E(*n*) Equivariant Graph Neural Network to generate graph embeddings; (2) a surface-oriented point cloud (middle, blue box) obtained from the entire protein is processed via a quasi-geodesic convolution layer to capture the geometric and chemical properties of the protein’s surface; and (3) the protein amino acid sequence (bottom, pink box) is processed by ESM2 to produce sequence-based embeddings that contain evolutionary and functional information about the protein. Together, these three featurization approaches encode complementary information for integration in T-ALPHA, enabling comprehensive modeling of the protein for downstream protein-ligand binding affinity prediction. The protein-ligand complex structure shown corresponds to PDB ID 3ACX.^110^

**Figure 2.**
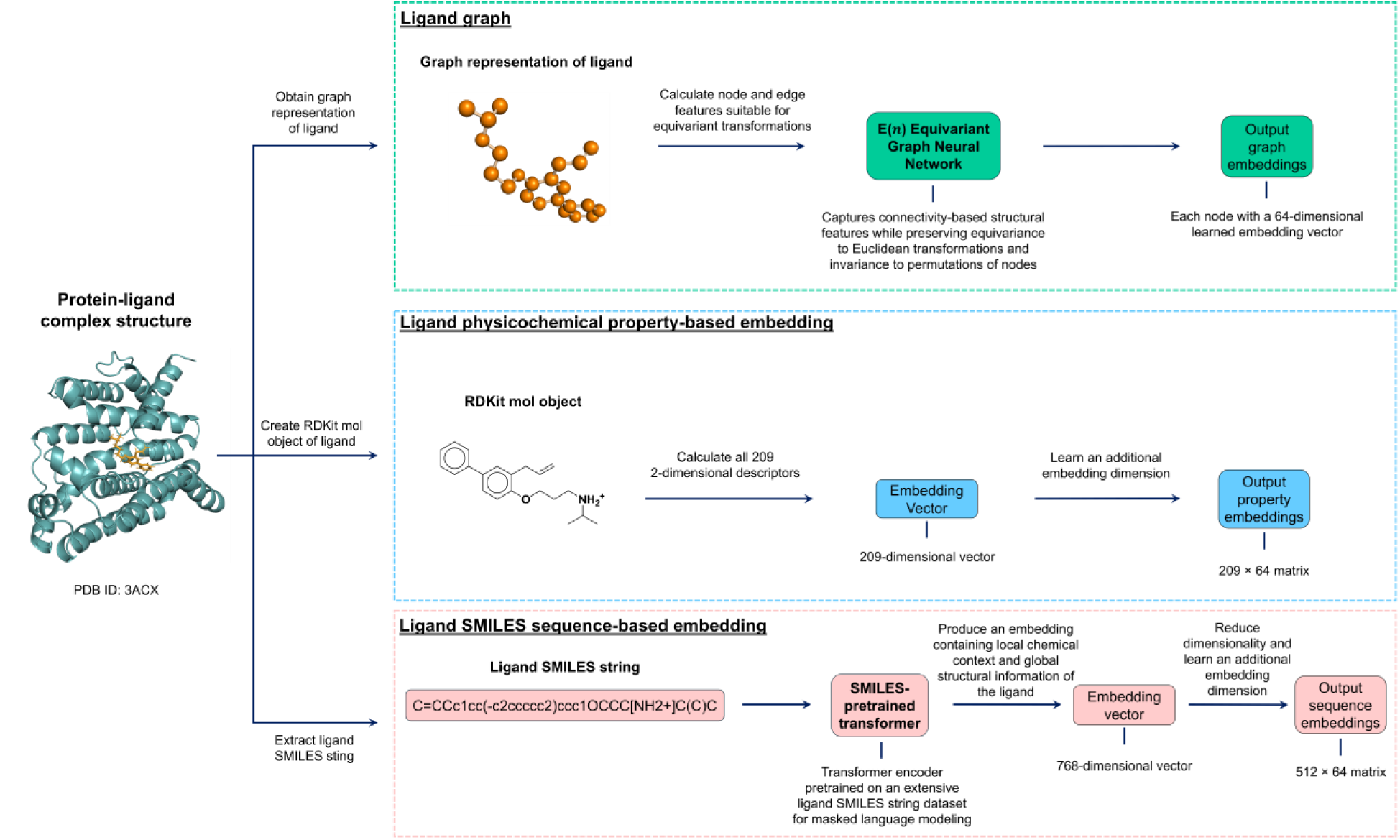
The ligand channel of the T-ALPHA pipeline. The ligand is represented and processed in three distinct ways: (1) a graph representation (top, green box), where nodes and edges are annotated with atomic and chemical features, is processed by an E(*n*) Equivariant Graph Neural Network to generate graph embeddings; (2) physicochemical property-based embeddings (middle, blue box) capture molecular-level features that contribute to binding affinity such as size, polarity and flexibility; and (3) the ligand SMILES string (bottom, pink box) is processed by a pretrained transformer to obtain embeddings that capture local substructures, stereochemistry, and long-range dependencies. Together, these three featurization approaches encode complementary information for integration in T-ALPHA, enabling comprehensive modeling of the ligand for downstream protein-ligand binding affinity prediction. The protein-ligand complex structure shown corresponds to PDB ID 3ACX.^110^

##### 2.3.1.1 Protein Pocket Graph

For the protein pocket graph representation, each non-hydrogen atom of the protein pocket is represented as a node, and each covalent bond between these atoms is represented as an edge. The nodes and edges of this graph are richly annotated with features that capture both atomic properties and chemical interactions that are suitable for equivariant transformations.

The node features are described in Table 1, and the edge features are described in Table 2.

**Table 1.**
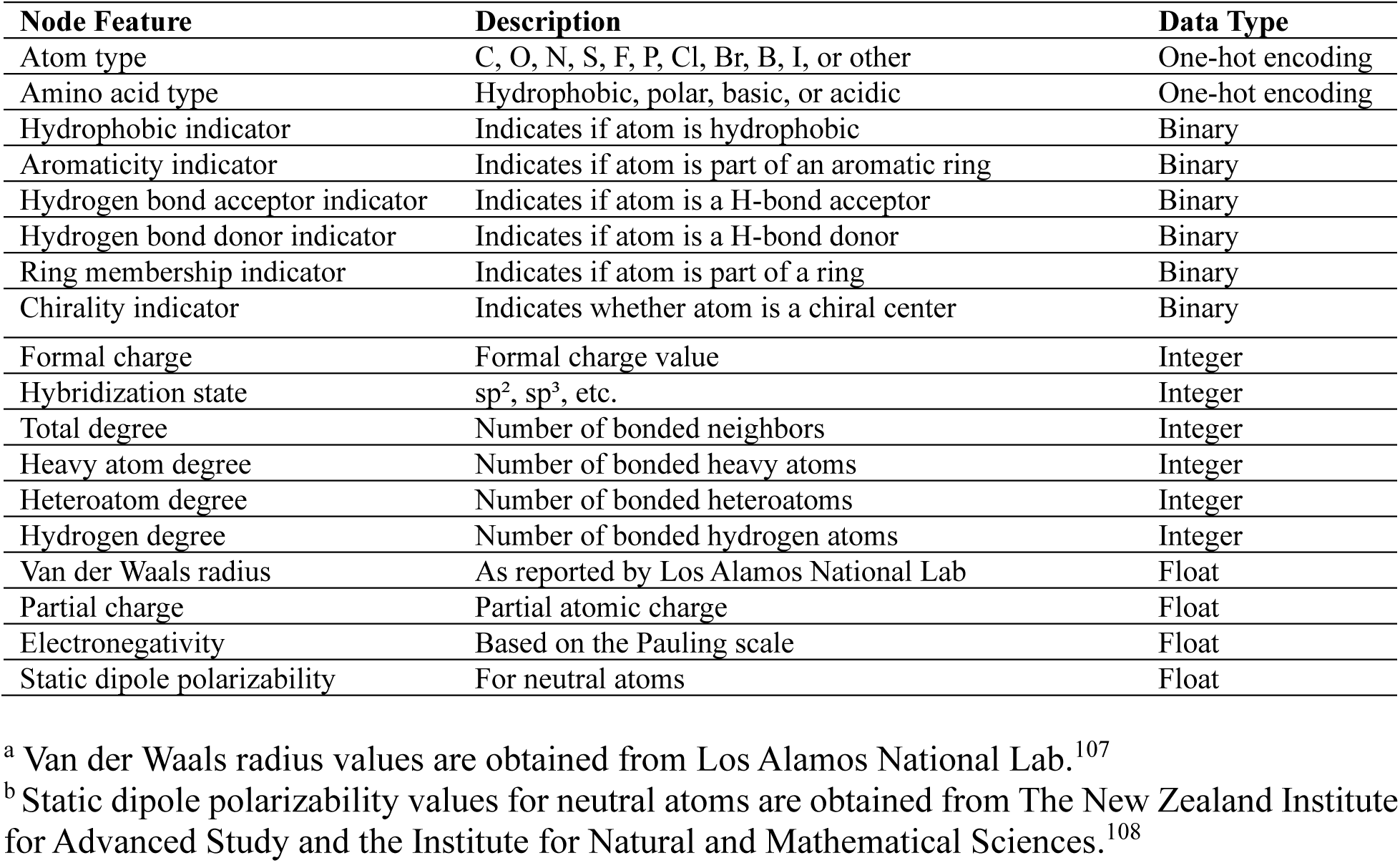
Node features used for protein pocket graph representation.

**Table 2.** Edge features used for protein pocket graph representation.

| Edge Feature | Description | Data Type |
| --- | --- | --- |
| Bond order | Interatomic bond order | Integer |
| Aromaticity indicator | Indicates if bond is part of an aromatic ring | Binary |
| Ring membership indicator | Indicates if bond is part of a ring | Binary |
| Interatomic distance | Measured in angstroms (Å) | Float |
| Electronegativity difference | Absolute difference in electronegativity between the two bonded atoms (based on the Pauling scale) | Float |
| Electrostatic interaction energy | Coulombic interatomic electrostatic interaction energy | Float |

<sup>a</sup> Electrostatic interaction energy is calculated as $q_i q_j / r_{ij}^2$ where $q_k$ denotes the partial charge of atom $k$ and $r_{kl}$ represents the interatomic distance (Å) between atoms $k$ and $l$ .

##### 2.3.1.2 Protein Pocket Surface-Based Point Cloud

In order to capture the detailed geometry and chemical properties of the protein-ligand interface, T-ALPHA employs an approach largely based on the dMaSIF^109^ method for describing the curvature of the protein pocket. This approach involves converting atomic-level information into an oriented point cloud that represents the surface of the protein pocket, and then processing this point cloud via quasi-geodesic convolutions. In our implementation, the point cloud is obtained from an disconnected atomic graph created identically to that described in Section 2.3.1.1 but with a few exceptions: (1) it is derived from the entire protein rather than just the protein pocket, (2) it also considers hydrogen atoms as nodes in addition to non-hydrogen atoms, and (3) it does not contain any edges.

This unconnected graph representation is processed through a multi-step pipeline designed to accurately capture the relevant protein pocket surface geometry and chemical characteristics for each protein-ligand complex. First, random points are placed around each atom of the protein, and their positions are iteratively refined using a soft distance metric which evaluates how close each sampled point is to the actual surface boundary, taking into account both the atomic radii and local geometric constraints. Through gradient-based optimization, these points converge to positions where the distance metric aligns with a defined threshold representing the protein’s surface. This results in a dense collection of points that accurately reflects the topography of the protein surface. Once these initial points have converged, a cubic grid clustering method is applied to produce a more uniform representation.^109^ Next, surface normals are computed for each remaining point, where the normal vectors are derived from the gradient of the soft distance function used to position the points on the protein surface. Each normal vector therefore indicates the outward-facing direction of the surface at the respective point, capturing orientation-based information for downstream transformations (see Section 3.1.2).

##### 2.3.1.3 Protein Amino Acid Sequence-Based Embedding

In addition to the information derived from the protein’s 3D structure, T-ALPHA also integrates embeddings obtained from ESM2 — a large-scale transformer model pretrained on extensive protein sequence databases.^111^ ESM2 processes each amino acid sequence using self-attention mechanisms to capture complex relationships and dependencies between amino acid residues, thereby encoding evolutionary information, structural context, and functional insights from the sequence. The output is a high-dimensional embedding vector that encapsulates relevant sequence-based information such as residue conservation, proximity to active sites, secondary structure elements, and overall fold constraints, rooted in the extensive evolutionary and structural patterns learned by ESM2 during its large-scale pretraining. By leveraging ESM2, we can extract complementary features that are not directly inferable from the protein pocket graph or the protein pocket surface-based point cloud. For example, certain amino acid residues might be critical for function and highly conserved evolutionarily, which ESM2 can highlight even if these residues do not have immediately obvious structural patterns.

#### 2.3.2 Ligand Featurization

For the ligand channel in T-ALPHA, we employ a featurization strategy analogous to that used for the protein channel, capturing multiple facets of the ligand’s chemical and structural properties that we later demonstrate are important for predicting protein-ligand binding affinity. Specifically, we represent and process the ligand in three distinct ways: (1) a sparse graph which is processed by an E(*n*) EGNN to learn connectivity-based structural features, (2) a molecular-level descriptor vector which is processed by an MLP to encapsulate global physicochemical properties, and (3) a SMILES sequence-based embedding which is further refined by an MLP to capture local and global chemical context, such as atom-level interactions, substructures, and stereochemical details, that are not inferable from the graph or global descriptor representations. This multifaceted approach ensures a comprehensive representation of the ligand for downstream tasks.

##### 2.3.2.1 Ligand Graph

The ligand is represented as a graph where each non-hydrogen atom is a node, and each covalent bond between these atoms is an edge. The node and edge features are calculated similarly to those in the protein pocket graph (Section 2.3.1.1), with the only distinction being that amino acid type is not one of the node features.

##### 2.3.2.2 Ligand Physicochemical Property-Based Embedding

In addition to the graph representation, we compute a molecular-level descriptor vector for each ligand using RDKit.^106^ This descriptor vector encapsulates a range of physicochemical and topological properties that provide a comprehensive overview of the ligand’s chemical characteristics. These descriptors include information on molecular size, polarity, flexibility, electronic properties, and other relevant features that influence binding affinity. By characterizing the ligand’s global properties, this descriptor vector complements the detailed local structural information captured by the graph representation, ensuring that both global and local features contribute to the overall ligand representation.

##### 2.3.2.3 Ligand SMILES Sequence-Based Embedding

As an additional feature representation of the ligand, we utilize a transformer encoder pretrained on a large dataset of SMILES strings to extract a context-rich embedding that contains information about complex chemical patterns and substructures. This pretrained model utilizes self-attention mechanisms to capture both local chemical contexts such as functional groups and ring systems, as well as global structural information.

In this approach, the ligand SMILES string is first tokenized into individual symbols representing atoms, bonds, and structural features. These tokens include elements like C, N, O; bond types denoted by symbols such as “=”, “#”; branching symbols like “(”, “)”; and stereochemical indicators such as “@”, “/”, “\”. Each token is mapped to a learnable high-dimensional embedding vector, and each positional index in the input is encoded into a learnable high-dimensional vector. The token embeddings capture semantic information about the chemical symbols, while the positional encodings provide the transformer with information about the order of the tokens in the sequence. This is critical because the self-attention mechanism in transformers is inherently permutation-invariant and does not consider token order unless explicitly encoded. The final input embeddings are obtained by element-wise summing the token embeddings and positional encodings, producing a combined representation for each token that integrates both semantic and positional information.

These combined embeddings are then passed through a series of transformer encoder blocks. Each block consists of a multi-head self-attention sublayer followed by a residual connection and layer normalization, and a position-wise feed-forward network sublayer which is also followed by a residual connection and layer normalization. The self-attention mechanism allows the model to compute a weighted representation of all tokens in the sequence relative to a given token, capturing intricate patterns and dependencies within the molecule. For each token, self-attention computes three vectors: a query vector, a key vector, and a value vector, all derived from learned linear transformations of the input embedding. The attention score between any two tokens is calculated as the scaled dot product of the query vector of one token with the key vector of another, normalized using a softmax function to ensure the scores sum to one. These scores are then used to compute a weighted sum of the value vectors, resulting in an output embedding that reflects the token’s context in the sequence. By attending to all tokens in the sequence, the model effectively captures both local interactions, such as adjacent atoms in a functional group, and long-range dependencies, such as conjugated systems or distant functional groups that influence each other. In the multi-head self-attention mechanism, multiple attention heads process the embeddings in parallel, with each head learning to focus on different types of relationships among tokens. This parallel processing enhances the model’s ability to capture diverse aspects of chemical interactions within the molecule.

Following the multi-head self-attention sublayer, each transformer block applies a residual connection and layer normalization to the output of the attention mechanism. Next, a position-wise feed-forward sublayer is applied to each token individually. This feed-forward sublayer consists of two linear transformations separated by a Gaussian Error Linear Unit (GELU) activation function. The feed-forward sublayer is followed by another residual connection and layer normalization. This setup enables the effective modeling of intricate patterns and long-range dependencies across sequences, with residual connections promoting stable gradient flow and layer normalization ensuring numerical stability to facilitate efficient training of deep architectures. The transformer that we use contains 10 sequential transformer blocks.

After the input SMILES string is processed by the transformer blocks, we obtain contextualized embeddings for each token in the sequence, where each embedding now contains information about the token itself and about its relationships with other tokens in the sequence. In our implementation, we extract the embedding corresponding to the start token added at the beginning of the sequence. This start token aggregates information from all positions in the sequence during the self-attention computations, effectively capturing a global representation of the molecule.

For each datapoint, the start token embedding extracted from the pretrained transformer is passed through an MLP to optimize it for predicting protein-ligand binding affinity. This step allows the model to refine the embedding, capturing subtle chemical features and intricate patterns that might not be inferable from the ligand’s graph representation or molecular-level descriptors.

#### 2.3.3 Protein-Ligand Complex Featurization

In T-ALPHA, the protein-ligand complex is represented as a unified graph that integrates both the protein pocket and ligand into a single structure, designed to capture the intricate interactions between the two molecules that are crucial for accurate binding affinity prediction.

The nodes in this graph represent non-hydrogen atoms from both the protein and the ligand. The node features are identical to those previously described for the protein (Section 2.3.1.1) and ligand (Section 2.3.2.1) graphs, with the addition of a source identifier that indicates whether a given node belongs to the protein or ligand, enabling the model to learn source-specific patterns and interactions.

Edges in the graph represent both intramolecular and intermolecular interactions. Intra-molecular edges are established based on covalent bonds within the protein and within the ligand, as defined in their respective individual graphs. These edges capture the internal connectivity of each molecule. Intermolecular edges are introduced between protein and ligand non-hydrogen atoms that are within 4.5 Å to capture potential non-covalent interactions critical for binding, such as hydrogen bonds, hydrophobic contacts, π-π stacking, and electrostatic interactions. We chose 4.5 Å as the distance threshold because it is commonly used as a hydrophobic distance threshold in protein-ligand interaction modeling.^112^ Edge features are identical to those used for the protein and ligand graphs, with an additional interaction feature to indicate whether a given edge represents an intramolecular connection or an intermolecular interaction. This labeling enables the model to distinguish between bonds that define molecular structures and interactions that contribute to binding affinity that are not apparent when considering the protein and ligand separately. By integrating the protein and ligand into a single graph with enriched node and edge features, T-ALPHA is able to consider both the structural details of the individual molecules and the critical interactions between them.

#### 2.3.4 Data Scaling

To ensure that our model effectively learns from features of varying scales and to enhance training stability, we apply standardization to all continuous features across our dataset. Specifically, we transform these features to have a mean of zero and a standard deviation of one. This preprocessing step is crucial for preventing any single feature from disproportionately influencing the learning process due to differences in magnitude.

Across our graph representations (i.e., those for the protein pocket, the entire protein used in the dMaSIF-based module, the ligand, and the protein-ligand complex), we standardize continuous node features including van der Waals radius, partial charge, electronegativity, and polarizability. For the edge features, we standardize interatomic distance, electronegativity difference, and Coulombic electrostatic interaction energy. In addition to the graph-based features, we standardize the protein amino acid sequence-based embeddings, ligand physicochemical property-based embeddings, and ligand SMILES sequence-based embeddings.

#### 2.3.5 Training and Validation Set Construction

For each training-validation-test split, the test set is predefined and excluded entirely from the training and validation data. To partition the remaining data, we use a structured and reproducible approach. For CASF 2016, 95% of the remaining PDBbind general set is randomly selected for training, while the remaining 5% is used for validation. For LP-PDBbind, the predefined splits provided by the authors are used directly to ensure consistency with previous studies. For the BDB2020+, Mpro, and EGFR test sets, a single model is trained with 95% of the remaining PDBbind general set randomly selected for training, while the remaining 5% is used for validation.

## 3 Architecture

T-ALPHA employs a hierarchical multimodal transformer-based architecture designed to integrate multiple complementary feature representations from three channels (protein, ligand, and protein-ligand complex) to accurately predicting binding affinity.

The protein channel contains three architectural components: (1) an E(*n*) EGNN that processes a graph representation of the protein pocket, (2) an adaptation of the dMaSIF^109^ model based on quasi-geodesic convolutions that processes a point cloud representation of the protein surface, and (3) an MLP that processes the protein amino acid sequence-based embeddings. The output of the dMaSIF-based model is passed to a transformer encoder, and the outputs of the E(*n*) EGNN and the MLP are passed to separate transformer decoders. Each of the decoders engages in cross-attention with the encoder output, enabling the integration of geometric features of the protein pocket surface into both the graph-based connectivity features and the sequence-based evolutionary and functional features. The outputs from each of the three transformers are then concatenated and reduced in dimensionality to match the output dimensionality of the protein-ligand complex E(*n*) EGNN, ultimately resulting in a single embedding to describe the protein channel.

For the ligand channel, an analogous architectural framework is employed, containing three components tailored specifically to capture the multifaceted characteristics of the ligand: (1) an E(*n*) EGNN that processes a graph representation of the ligand, (2) an MLP that processes the ligand physicochemical property-based embeddings, and (3) an MLP that processes the ligand SMILES sequence-based embeddings. The output of the MLP that processes the property-based embeddings is passed to a transformer encoder, and the outputs of the E(*n*) EGNN and the MLP that processes the sequence-based embeddings are passed to separate transformer decoders. Each decoder engages in cross-attention with the encoder output, enabling the integration of molecular-level physicochemical features into both the graph-based connectivity features and the sequence-based chemical and structural features. Similarly to as is done for the protein channel, the outputs from each of the three transformers are then concatenated and reduced in dimensionality to match the output dimensionality of the protein-ligand complex E(*n*) EGNN, ultimately resulting in a single embedding to describe the ligand channel.

The protein-ligand complex channel focuses exclusively on processing the graph of the bound complex using an E(*n*) EGNN. This approach captures the spatial relationships and interactions between the protein and ligand atoms that are important for accurately predicting binding affinity.

The outputs from all three channels are integrated using the proposed hierarchical transformer framework (Figure 3). The output of the protein-ligand complex channel is passed to a transformer encoder, and the outputs of the protein and ligand channels are passed to separate transformer decoders. Each decoder engages in cross-attention with the encoder output, enabling the integration of detailed binding interaction information from the protein-ligand complex with the rich representations of the individual protein and ligand. The outputs from the three transformer layers are then concatenated, pooled with an attention mechanism, and passed to a final MLP to produce the output prediction.

**Figure 3.**
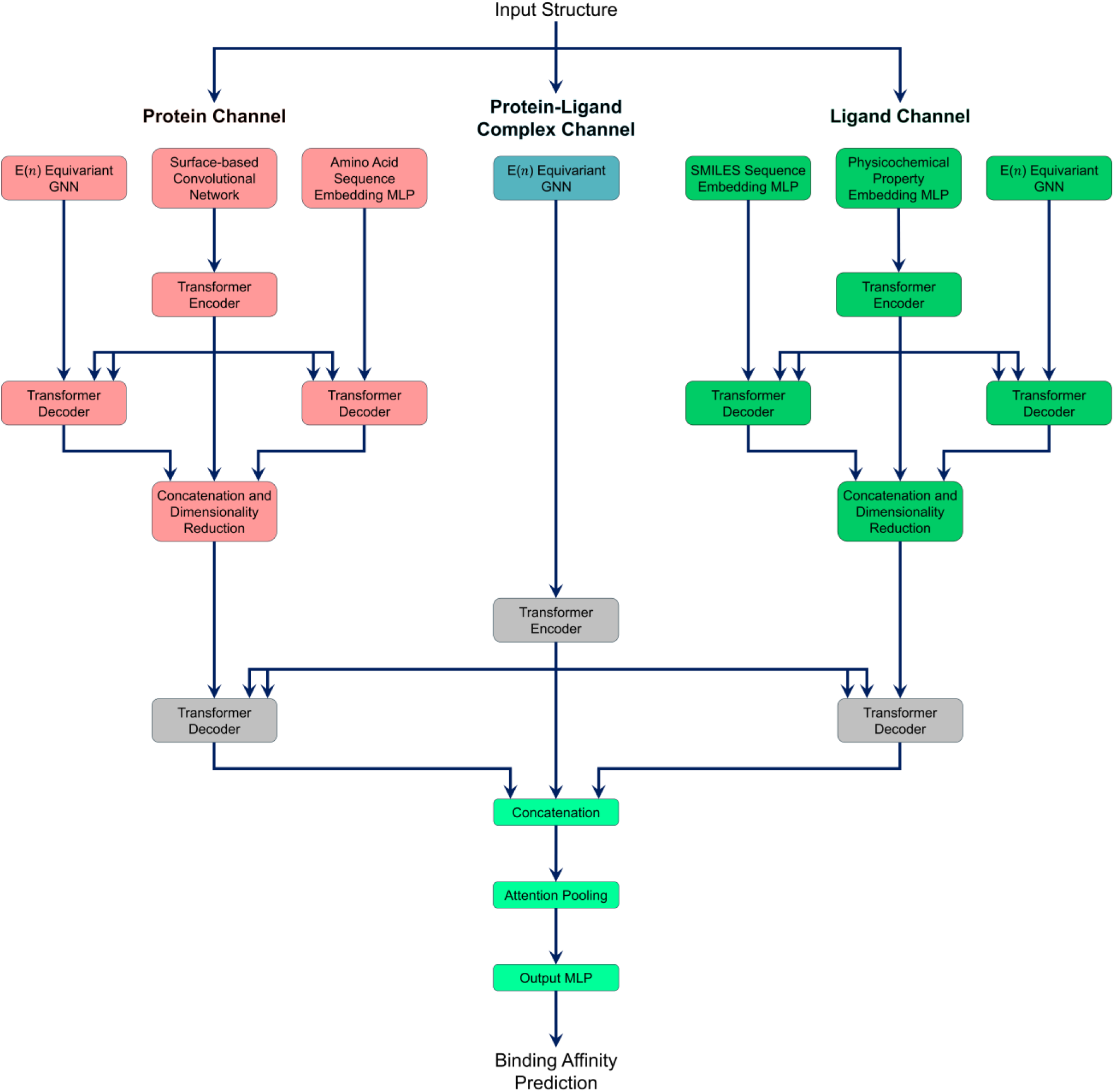
Overview of the hierarchical multimodal transformer-based architecture of T-ALPHA for protein-ligand binding affinity prediction. A given input protein-ligand complex structure is first processed by protein, ligand, and protein-ligand complex channels. The outputs of each of the three channels are then further processed by transformers, the outputs of which are then concatenated, pooled with an attention-based method, and passed to a final MLP to generate the output prediction.

### 3.1 Protein Channel

#### 3.1.1 E(*n*) Equivariant Graph Neural Network

To capture the connectivity-based structural features of the protein pocket while preserving E(*n*) equivariance (i.e., equivariance under Euclidean transformations including rotations, translations, reflections, and permutations within n-dimensional Euclidean space), we employ an adaptation of the E(*n*) EGNN presented by Satorras et al.^113^ This architectural component ensures that the learned graph representation accurately reflects the spatial relationships and geometric structure of the protein pocket, enabling the model to capture critical interactions and patterns that are independent of the molecular orientation or position.

In our E(*n*) EGNN, each node *i* represents a non-hydrogen atom in the protein pocket, characterized by initial features **h**_*i*_ that encode atomic properties (Table 1), and coordinates **x**_*i*_ representing the atom’s 3D position. Edges (*i*, *j*) connect covalently bonded atoms and include edge features **e**_*ij*_ (Table 2).

The E(*n*) EGNN updates both the node features and coordinates through iterative message passing, where messages (**m**_*ij*_) are computed along the edges:

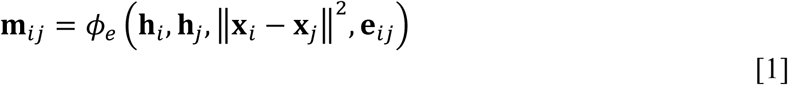

where ‖**x**_*i*_− **x**_*j*_‖ is the squared Euclidean distance between nodes *i* and *j*, and Φ*e* is an MLP that computes edge-specific messages **m**_*ij*_ by integrating node features, edge features, and geometric distance. In addition, we apply a learned attention mechanism that assigns a weight to each edge, allowing the model to focus on more important interactions. For the protein pocket graph, each atom’s feature vector **h**_*i*_ has a dimensionality of 31, and each edge feature vector **e**_*ij*_ has a dimensionality of 6.

The node coordinates are updated based on the messages:

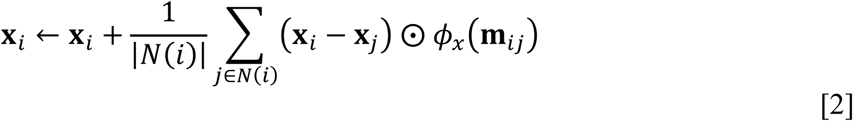

where *N*(*i*) denotes the neighbors of node *i*, *Φ*_*x*_ is a coordinate-based MLP, and ⨀ denotes element-wise multiplication. This update mechanism ensures that coordinate transformations depend on relative positions and learned messages, thus maintaining E(*n*) equivariance. We aggregate the coordinate updates by taking the mean over the neighbors.

The node features are updated using the aggregated messages:

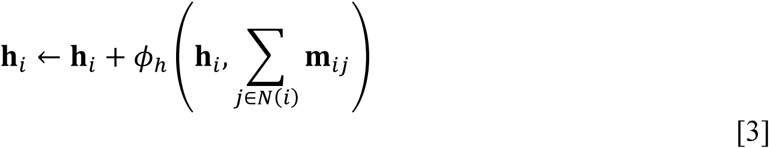

where *Φ*_ℎ_ is an MLP that integrates the incoming messages to refine the node features. We incorporate residual connections by adding the original node features **h**_*i*_ to the output of *Φ*_ℎ_, thus stabilizing the training.

The protein pocket E(*n*) EGNN consists of four layers, with each layer applying updates to the node features and coordinates to progressively refine the protein pocket graph representation. Each node’s final feature vector has a dimensionality of 64, capturing the spatial and relational information necessary for accurately predicting protein-ligand binding affinity.

#### 3.1.2 Quasi-Geodesic Convolutional Layer

To capture the detailed geometry and curvature of the protein pocket surface, we employ a dMaSIF^109^–based quasi geodesic convolutional layer to process a surface-based point cloud representation of the protein pocket.

The protein pocket surface is represented as a point cloud of 512 points, where each point **x**_*i*_ ∈ ℝ^3^ lies on the molecular surface of the protein. For each point **x**_*i*_, we compute a smoothed normal vector **n**_*i*_ by averaging the normals of neighboring points **n**_*j*_ within a Gaussian kernel:

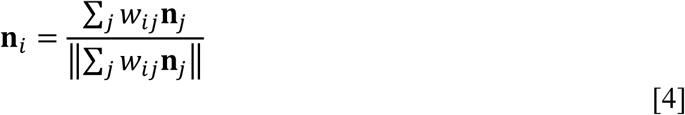

where each weight *w*_*ij*_ are defined as:

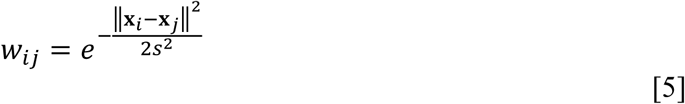

and *s* is a scale parameter controlling the spread of the Gaussian smoothing window. We implement five different scale parameter values: 1.0, 2.0, 3.0, 5.0, and 10.0, for computing five geometric features at each point. Using the smoothed normal vector **n**_*i*_, we compute two orthogonal tangent vectors **u**_*i*_ and **v**_*i*_ to form an orthonormal basis [**n**_*i*_, **u**_*i*_, **v**_*i*_] at each point, thus providing a local coordinate system at each point on the surface.

We estimate the mean curvature *H*_*i*_ and Gaussian curvature *K*_*i*_ at each scale for each point using a local quadratic approximation of the surface. Specifically, we compute the shape operator **S**_*i*_ by solving:

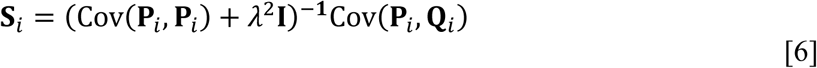

where **P**_*i*_ and **Q**_*i*_ are matrices of coordinate differences and normal differences, respectively, that have been projected into their local tangent planes, *λ* is a regularization parameter which is set to 0.1 Å, and **I** is the identity matrix. Cov(⋅, ⋅) is an element-wise-weighted covariance matrix where the weights are derived from the Gaussian smoothing window. The mean and Gaussian curvatures are then given by:

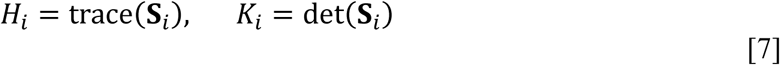

In addition to geometric features, we incorporate chemical information by employing a separate module with message passing, where chemical properties of atoms in proximity to a given surface point are propagated to that point. Each atom *a*_*j*_ is associated with a feature vector **f**_*j*_ encoding its chemical properties. In our implementation, each atom’s feature vector has a dimensionality of 32, incorporating the features listed in Table 1 with the addition of a one-hot encoding for the Hydrogen atom type. We apply a neural network *Φ*_*f*_ to transform these features:

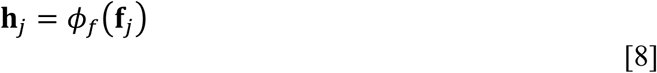

We then perform message passing among atoms to update their features based on their 16 nearest neighboring atoms:

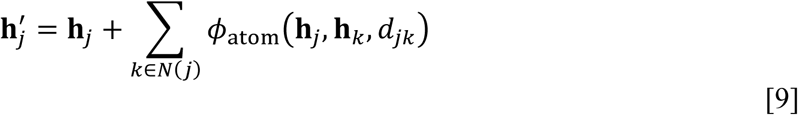

where *Φ*_atom_ is a neural network function, *N*(*j*) denotes the 16 nearest neighboring atoms to atom *j*, and *d*_*jk*_ is the distance between atoms *j* and *k*. We use three message passing layers to iteratively update the atomic features.

We then propagate the updated atomic features to each surface point by considering the 16 nearest neighboring atoms:

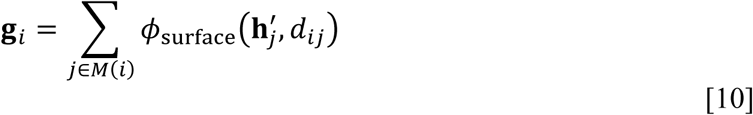

where *Φ*_surface_ is a neural network function, *M*(*i*) denotes the 16 nearest atoms to surface point *i*, and *d*_*ij*_ is the distance between surface point *i* and atom *j*. The resulting chemical feature vector **g**_*i*_, which has a dimensionality 32, is concatenated with the geometric features (i.e., the mean and Gaussian curvatures computed at five scales). This results in a combined feature vector **c**_*j*_ with a dimensionality of 42.

To aggregate these combined geometric and chemical features over the surface, we apply a quasi-geodesic convolutional layer that updates the combined feature vector **c**_*j*_ at each point by aggregating information from neighboring points using a learned kernel:

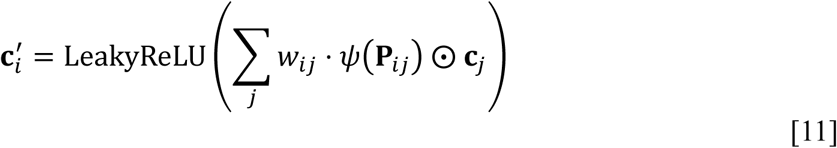

where *w*_*ij*_ is a scalar weight based on a pseudo-geodesic distance, and *ψ* is a neural network acting on the local coordinate differences **P**_*ij*_. The weighting function used to obtain *w*_*ij*_ considers both spatial proximity and normal similarity:

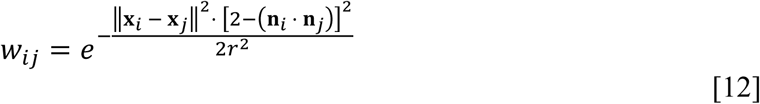

where *r* is a scale parameter that controls the sensitivity of the weighting function to spatial proximity and normal similarity, which we set to 9.0 Å.

For each of the 512 surface points, the final embedding **c**^′^ is a vector of dimensionality 64 that encapsulates rich geometric and chemical characteristics of the protein pocket surface.

#### 3.1.3 Amino Acid Sequence-Based Embedding Neural Network

To incorporate global evolutionary and biophysical characteristics of the protein that can be derived from its amino acid sequence, we utilize ESM2, a protein language model trained for masked token prediction, to produce a fixed-dimensional embedding vector **e**_seq_ ∈ ℝ^2560^. We reduce the dimensionality of each embedding from 2_560 to 512 using a linear layer, followed by batch normalization and ReLU activation:

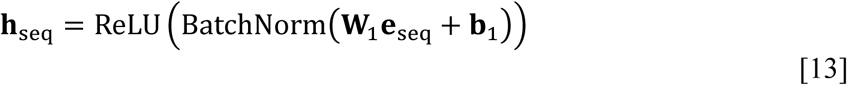

where **W**_1_ ∈ ℝ^512×2560^ is a weight matrix, and **b**_1_ ∈ ℝ^512^ is a bias vector. The resulting vector **h**_seq_ ∈ ℝ^512^ is then reshaped into a sequence by treating each of its 512 elements as individual tokens with scalar features:

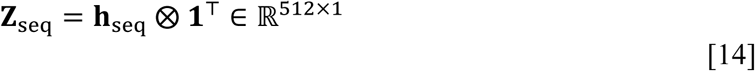

where ⨂ denotes the tensor product operation and **1** is a vector of ones. We then expand the feature dimension of each token of the resulting vector **Z**_seq_ ∈ ℝ^512×1^ from 1 to 64 using another linear layer, where:

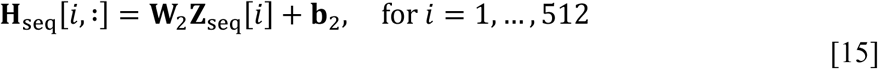

where **W**_2_ ∈ ℝ^64×1^, and **b**_2_ ∈ ℝ^64^. The resulting representation **H**_seq_ ∈ ℝ^512×64^ is now suitable for processing by a transformer decoder.

#### 3.1.4 Protein Transformer

To effectively integrate the diverse protein representations obtained from the E(*n*) EGNN, the quasi-geodesic convolutional layer, and the amino acid sequence-based embedding neural network, we employ a protein transformer architecture composed of a transformer encoder and two transformer decoders, enabling different feature modalities to interact through cross-attention mechanisms. The key technical difference between a transformer encoder and decoder lies in the use of a causal mask in the decoder’s self-attention mechanism. This mask ensures that, at each decoding step, the model only attends to current and previous tokens. In contrast, the encoder processes all input tokens simultaneously without such restrictions.

The processed surface features from the dMaSIF-based model, denoted as **H**_surf_ ∈ ℝ^512×64^ (where 512 is the number of surface points and 64 is the feature dimensionality), are passed through a transformer encoder to capture contextual relationships among the learned surface point embeddings:

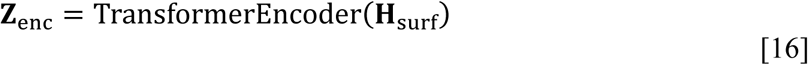

The outputs of the E(*n*) EGNN (**H**_graph_ ∈ ℝ^*N* nodes×64^) and the amino acid sequence-based embedding neural network (**H**_seq_∈ ℝ^512×64^) are each passed to separate transformer decoders, each of which incorporates the transformer encoder output via cross-attention:

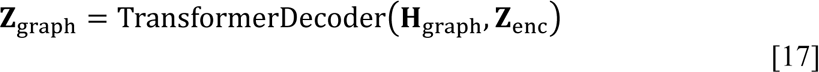

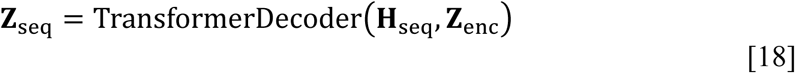

The cross-attention mechanism allows the transformer decoders to selectively focus on relevant parts of the encoder’s output. Specifically, for a decoder input **H**_dec_(either **H**_graph_ or **H**_seq_), cross-attention is computed as:

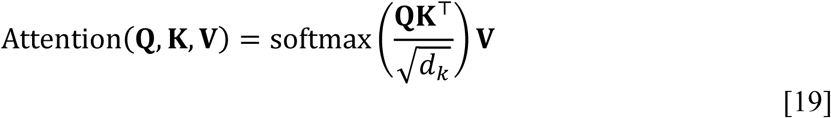

where queries **Q** = **H**_dec_**W**_*Q*_, keys **K** = **Z**_enc_**W**_*K*_, values **V** = **Z**_enc_**W**_*V*_, and **W**_*Q*_, **W**_*K*_, and **W**_*V*_ are learned projection matrices. The dimensionality of each of the key vectors, *d*_*k*_, is used to scale the dot products.

The graph decoder output **Z**_graph_ for each protein has a uniform length within the batch during decoding, achieved by padding all graphs to match the number of nodes in the largest graph in the batch. After decoding, the padding is removed, resulting in an effective length of **Z**_graph_ that matches the number of nodes in the respective graph. To obtain a fixed size representation for each protein graph, we apply a masked mean pooling over the node dimension:

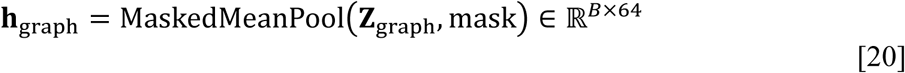

where the mask accounts for the variable node lengths in the batch. Here, *B* denotes the batch size. The pooled graph representation **h**_graph_ ∈ ℝ^*B*×64^ is reshaped and expanded via a learned linear layer to match the dimensionality of the other modalities, resulting in **Z**′_graph_ ∈ ℝ^512×64^.

We then concatenate the encoder output **Z**_enc_, the processed graph decoder output **Z**′_graph_, and the sequence decoder output **Z**_seq_ along the sequence length dimension to obtain a unified representation of the protein:

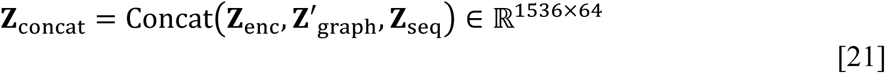

We permute the dimensions of **Z**_concat_ to obtain **Z**^′^_concat_ ∈ ℝ^*B*×64×1536^, and then apply a linear projection to reduce the concatenated feature dimension from 1_536 to 512. After permuting back to the original dimensions, we obtain the final protein representation **Z**^′^_protein_ ∈ ℝ^512×64^. This representation encapsulates the integrated information from all three modalities, and is prepared for downstream processing.

### 3.2 Ligand Channel

#### 3.2.1 E(*n*) Equivariant Graph Neural Network

To capture the structural and connectivity-based characteristics of the ligand, we represent each ligand as a graph and process it using the same E(*n*) EGNN architecture described in Section 3.1.1. The only distinction between the ligand graph and the protein pocket graph is that the ligand graph has one less node feature, corresponding to the amino acid indicator in the protein pocket graph.

#### 3.2.2 Physicochemical Property-Based Embedding Neural Network

To incorporate molecular-level physicochemical properties of the ligand, we compute a vector of descriptors based on the 2D molecular structure using RDKit, resulting in a fixed-length vector **d**_prop_ ∈ ℝ^209^. We reshape and expand the vector using a learned linear layer, where:

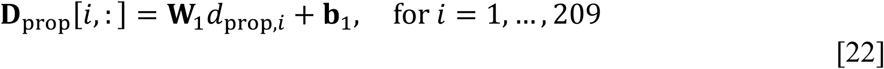

with **W**_1_ ∈ ℝ^64×1^ and **b**_1_ ∈ ℝ^64^. The resulting output **D**_prop_ ∈ ℝ^209×64^.

#### 3.2.3 SMILES Sequence-Based Embedding Neural Network

To capture complementary structural and chemical features of the ligand, we utilize a transformer encoder pretrained on a large dataset of SMILES strings to extract a contextual embedding **e**_seq_ ∈ ℝ^768^. To integrate these embeddings into our architecture, we reduce the embedding dimensionality from 768 to 512:

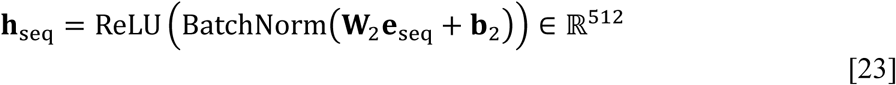

where **W**_2_ ∈ ℝ^512×768^ and **b**_2_ ∈ ℝ^512^. The resulting vector **h**_seq_ is then reshaped into a sequence of length 512, with each element representing a token, and its feature dimension is expanded using a learned linear layer:

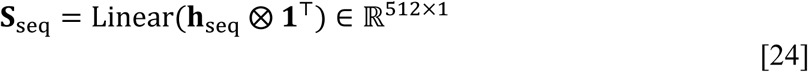

Each element of the reshaped sequence **S**_seq_ is projected into a higher-dimensional feature space using a learned linear transformation:

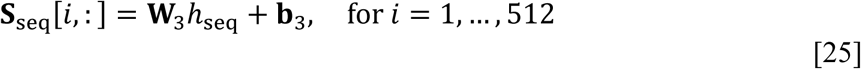

with **W**_3_ ∈ ℝ^64×1^ and **b**_3_ ∈ ℝ^64^.

#### 3.2.4 Ligand Transformer

To integrate the various ligand representations, we employ a transformer architecture analogous to that described in Section 3.1.4. The output of the physicochemical property-based embedding neural network serves as input to a transformer encoder. The outputs from the E(*n*) EGNN and the SMILES sequence-based embedding neural network are input into separate transformer decoders that incorporate the encoder output via cross-attention. The outputs from the three transformer components are then combined in the same manner as described for the protein transformer in Section 3.1.4. This combined ligand representation encapsulates integrated structural, physicochemical, and interaction-relevant substructural features.

### 3.3 Complex E(*n*) Equivariant Graph Neural Network

To model the critical interactions between the protein and ligand, we construct a unified graph representation of the protein-ligand complex and process it using an E(*n*) EGNN, as described previously in Section 3.1.1. An important distinction is that the protein and ligand E(*n*) EGNNs are each four layers, while the protein-ligand complex EGNN is eight layers. This increased depth allows the protein-ligand complex E(*n*) EGNN to capture intricate interactions between the protein and ligand atoms which are essential for accurately predicting binding affinity.

### 3.4 Meta Transformer

The output feature representations from each of the three channels are integrated via a meta transformer architecture that leverages cross-attention mechanisms to capture interdependencies among the protein, ligand, and complex that are critical for accurately predicting binding affinity. The output of the protein-ligand complex channel serves as the input to a transformer encoder. The outputs of the protein and ligand channels are each passed to separate transformer decoders, each of which uses the encoder output as the memory input in a cross-attention mechanism, allowing the protein and ligand modalities to attend to relevant parts of the complex encoding. The outputs of the protein-ligand complex encoder (**Z**_complex_), protein decoder (**Z**_protein_), and ligand decoder (**Z**_ligand_) are then concatenated along the sequence dimension to create a unified representation that aggregates the learned features from all three modalities:

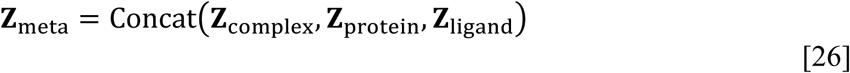

where **Z**_meta_ ∈ ℝ^1536×64^. Attention pooling is then applied to distill the concatenated representation into a fixed-size vector **v**_meta_ that captures the most relevant information across the sequence:

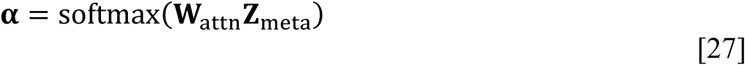

where **W**_proj_ projects the weighted sum of features to a fixed-size output vector **v**_meta_ ∈ ℝ^512^, and the attention weights **α** ∈ ℝ^1536×1^ are calculated as:

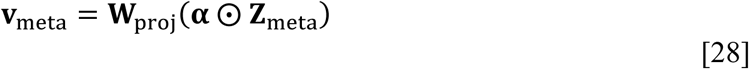

where **W**_attn_ is a learnable weight matrix for the attention mechanism. The pooled representation **v**_meta_ is then passed to an MLP to produce the final binding affinity prediction.

## 4 Model Training Details

### 4.1 Experimental Dynamic Range-Aware Custom Loss Function

The data from PDBbind contains target values *y*_*i*_, each associated with an operator *o*_*i*_ from the set {=, ∼, >, <}, indicating exact equality, approximate equality, greater than, or less than relationships, respectively. To account for these relational constraints, we designed a custom loss function ℒ that adjusts the penalization based on the operator associated with each target value. The loss for a given sample is defined as:

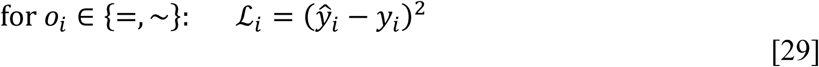

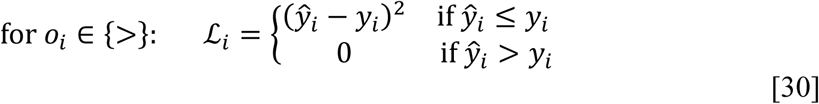

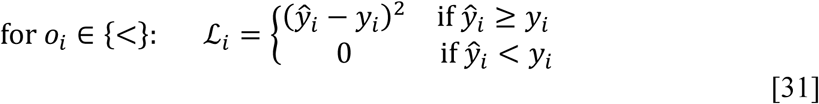

where 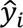 is the model’s prediction for the *i*-th sample. The overall loss is computed as the mean of the individual ℒ_*i*_ values over the batch:

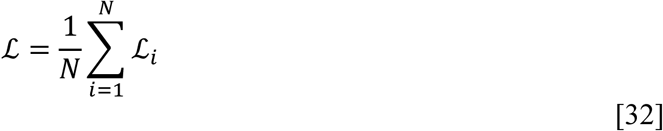

where *N* is the number of samples in the batch. This custom loss function ensures that the model respects the dynamic ranges of the experiments used to obtain binding affinity measurements, rather than assuming all labels to represent exact equalities.

### 4.2 Parameter Optimization

Optimization was performed using the AdamW optimizer,^114^ with an initial learning rate *n*_initial_ = 3 × 10^−4^ and a weight decay coefficient of 1 × 10^−5^. Gradient clipping with a maximum value of 0.1 was applied to ensure training stability.

We used an adaptive learning rate scheduler comprising two phases. The first phase is a warm-up period over the first *T*_warmup_ = 30 epochs, during which the learning rate increases linearly from 0.1 × *n*_initial_ to the initial learning rate *n*_initial_. The learning rate at epoch *t* during the warm-up phase is given by:

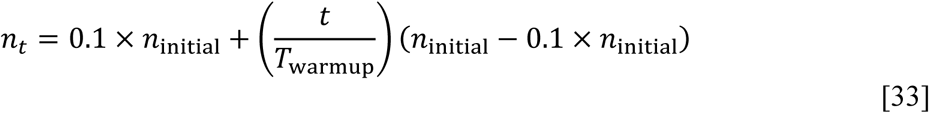

Following the warm-up phase, the learning rate is adjusted using a cosine annealing schedule over the remaining epochs, decreasing the learning rate from *n*_initial_ to a minimum learning rate *n*_min_ = 3 × 10^−5^ following a cosine curve:

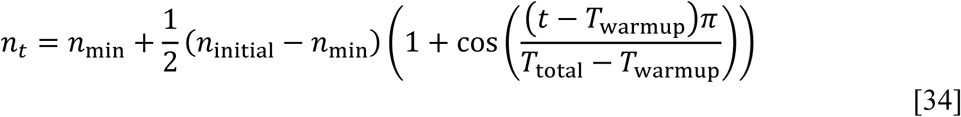

where the total number of training epochs *T*_total_ = 120.

### 4.3 Distributed Training and Scaling

We leveraged the Fully Sharded Data Parallel (FSDP) strategy implemented in PyTorch Lightning^115^ to distribute training across four NVIDIA A100 GPUs. This approach significantly reduces memory consumption and scales the training process efficiently, enabling an effective batch size of 128 (batch size of 32 per GPU).

### 4.4 Model Selection

We select the model parameters corresponding to the epoch with the lowest validation loss to be used inference. Learning curves for each of the trainings performed in this work are provided in Figures S1 – S3 in the *Supporting Information*.

## 5 Self-Learning Method for Protein-Specific Alignment

To improve ranking ability of compounds by binding affinity for specific protein targets without requiring additional experimental data, we propose a self-learning method that leverages uncertainty estimation and chemical similarity. We apply this approach to two protein targets, Mpro and EGFR, each associated with a set of ligands for which binding affinity data is available. These datasets serve as test sets for evaluating the proposed method.

The manifold hypothesis posits that high-dimensional data lies on low-dimensional manifolds embedded within the higher-dimensional space.^117^ We hypothesize that structurally similar compounds are expected to cluster together on these manifolds, sharing relevant properties that contribute to protein-ligand binding affinity. To exploit this hypothesis, we select compounds with similar ECFP4s to those in the test set, thereby constructing a pseudo-training set focused on the specific regions of the chemical manifold relevant to the test set.

Using Monte Carlo dropout, we estimate the uncertainty of model predictions for these pseudo-training compounds and then update the model parameters using a weighted loss function that emphasizes low-uncertainty pseudo-labels. This approach successfully aligns the model to relevant regions of the chemical manifold, enhancing its ability to rank test set compounds by binding affinity for both Mpro and EGFR (Section 6.4).

### 5.1 Bayesian and Statistical Learning Theoretical Inspiration

Our method is inspired by foundational principles from Bayesian inference and statistical learning theory. In Bayesian statistics, when dealing with observations of varying uncertainty, each observation should be weighted according to its precision (inverse variance) during parameter estimation. Given a pseudo-labeled input-output pair (*x*_*i*_, *y*_*i*_) with associated predictive uncertainty 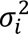, the likelihood function for the datapoint can be calculated as:

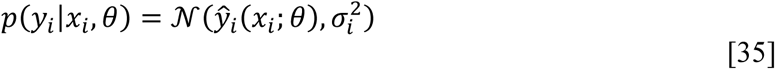

where *p*(*y*_*i*_|*x*_*i*_, *θ*) represents the probability of observing *y*_*i*_ given the input *x*_*i*_ and the model parameters *θ*. This probability is modeled as a Gaussian distribution *N* where the model’s prediction 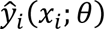 is the mean and the predictive uncertainty 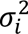 is the variance.

The negative log-likelihood over the dataset *D* composed of *M* points is:

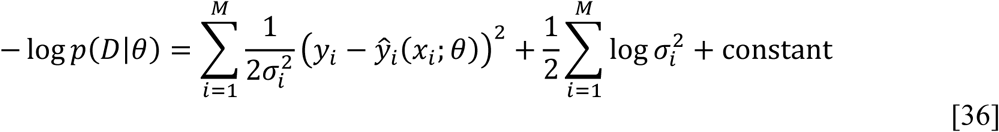

where *p*(*D*|*θ*) represents the joint likelihood of all observations in *D* given the model parameters *θ*. Assuming 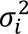 are known and fixed, and ignoring constant terms, the loss function simplifies to a precision-weighted mean squared error:

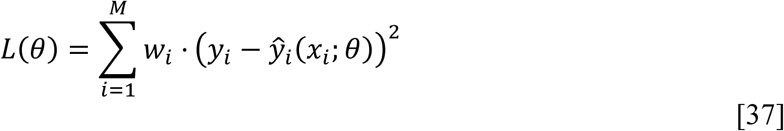

This weighting scheme ensures that datapoints with lower uncertainty have greater influence on the loss, thereby contributing more significantly to parameter updates.

Monte Carlo dropout approximates Bayesian inference in neural networks by interpreting dropout as a variational approximation to the posterior distribution *P*(*θ*|*D*).^116^ For each data point *x*_*i*_, the predictive mean 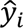 and variance 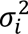 are estimated as:

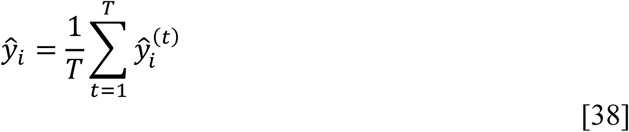

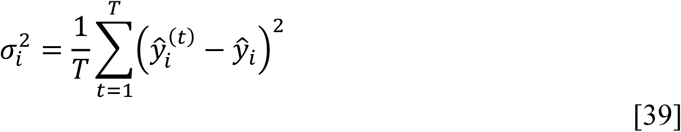

where *T* = 100 is the number of stochastic forward passes and 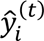 is the model prediction at iteration *t*.

To implement a numerically stable approximation of the precision-weighted loss (Equation 37), we adopt a smoothed weighting scheme that transforms variance values 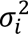 into weights to be incorporated into the loss function during parameter optimization:

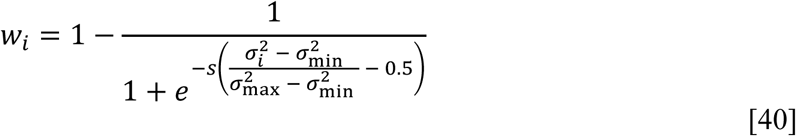

where 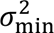 and 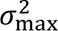 are the minimum and maximum variances in the pseudo-training dataset, respectively, and *s* = 10 is a scaling factor. This transformation ensures that weights smoothly decrease with increasing variance, promoting stability and preventing any single datapoint from disproportionately influencing the optimization process. (Figure S4 in the *Supporting Information*).

### 5.2 Implementation Details

For each of the test ligands for a given protein target, we compute the Tanimoto similarity between the corresponding Extended Connectivity Fingerprint with a diameter of 4 (ECFP4) and those of compounds in the pretraining set (∼ 5 million entries) used for the SMILES-based transformer encoder (Section 2.3.2.3). The Tanimoto coefficient *T* between two ECFP4s *A* and *B* is calculated as:

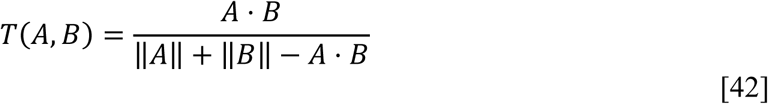

where *A* ⋅ *B* denotes the dot product of the binary vectors, and ‖*C*‖ represents the sum of the elements in *C*. For each test set ligand, we select the 10 compounds from the pretraining set with the highest Tanimoto similarity, resulting in a pseudo-training set of *N*^′^ = 10 × *N* compounds, where *N* is the number of datapoints in the test set.

For each pseudo-training compound, we generate a protein-ligand complex structure using Chai1 (see Section 9 for Chai1 implementation details). We then compute the pseudo-label and uncertainty-based weight for each pseudo-training datapoint as described in Section 5.1.

### 5.3 Experimental Setup and Validation

We implemented the self-learning method via two training strategies: fine-tuning a pre-trained T-ALPHA model and training the T-ALPHA architecture from scratch. Fine-tuning is performed with a learning rate of 3 × 10^−5^ for 200 epochs to ensure convergence. When training from scratch, we utilize the same procedure described in Sections 4.2 and 4.3, with the distinction that the model is trained for 200 epochs.

To validate the method, we performed control experiments where the uncertainty-based weights *w*_*i*_ were excluded from the loss function (Equation 37), allowing us to directly assess the contribution of uncertainty-based weighting to improved ranking performance on test set compounds.

Model parameters for inference are selected based on validation performance. Specifically, a validation set is created using the original *N* test compounds. Pseudo-labels and weights are computed for each validation datapoint using the procedures described in Section 5.1. The model’s performance is then evaluated on this validation set using the weighted loss function (Equation 37), and the parameters from the epoch achieving the lowest validation loss are chosen for inference.

## 6 Results

T-ALPHA was evaluated across multiple benchmarks to demonstrate its robustness in predicting protein-ligand binding affinity. We benchmarked on the CASF 2016 test set, widely regarded as the standard dataset for protein-ligand binding affinity scoring function assessment. While its popularity enables straightforward comparisons to existing models, we acknowledge the dataset’s flaws, such as significant data leakage between training and test sets, which artificially inflates performance metrics. To address these shortcomings, we also benchmarked T-ALPHA on datasets designed to mitigate data leakage and improve generalizability assessments: LP-PDBbind and BDB2020+ test sets. Additionally, we evaluated T-ALPHA on two protein-specific test sets corresponding to Mpro and EGFR to assess its applicability in scenarios that require high accuracy for specific protein targets.

### 6.1 Comparative Performance on the CASF 2016 Benchmark

T-ALPHA was benchmarked on the CASF 2016 test set and demonstrated superior performance across all evaluated metrics compared to every model reported in the literature to date (Table 3). T-ALPHA achieves the lowest Root Mean Square Error (RMSE: 1.112), the lowest Mean Absolute Error (MAE: 0.875), the highest Pearson correlation coefficient (*r*: 0.869), the highest coefficient of determination (*r*^2^: 0.738), and the highest Spearman rank correlation coefficient (*ρ*: 0.860). These results establish T-ALPHA as the current state-of-the-art deep learning model for predicting protein-ligand binding affinity (Figure S5 in the *Supporting Information*).

**Table 3.** Performance of T-ALPHA and the highest-performing models in the literature on the CASF 2016 benchmark.

| Model | RMSE | MAE | $r^2$ | Pearson $r$ | Spearman $\rho$ |
| --- | --- | --- | --- | --- | --- |
| T-ALPHA | **1.112** | **0.875** | **0.738** | **0.869** | **0.860** |
| T-ALPHA<sup>†</sup> | 1.134 | 0.893 | 0.721 | 0.857 | 0.843 |
| EHIGN<sup>51</sup> | 1.150 | N/R | N/R | 0.854 | N/R |
| TopoFormer-Seq<sup>60</sup> | 1.151 | N.R | N/R | 0.864 | N/R |
| MFE<sup>73</sup> | 1.151 | 0.882 | N/R | 0.851 | N/R |
| LGN<sup>54</sup> | 1.177 | 0.936 | N/R | 0.842 | N/R |
| GIGN<sup>39</sup> | 1.190 | N/R | N/R | 0.840 | N/R |
| HAC-Net<sup>33</sup> | 1.205 | 0.971 | 0.692 | 0.846 | 0.843 |
| TopBP<sup>119</sup> | 1.210 | N/R | N/R | 0.861 | N/R |
| CurvAGN<sup>45</sup> | 1.217 | 0.930 | N/R | 0.8305 | N/R |
| AEscore<sup>120</sup> | 1.22 | N/R | N/R | 0.83 | 0.64 |
| AK-score<sup>32</sup> | 1.22 | N/R | N/R | 0.812 | 0.670 |
| PLANET<sup>38</sup> | 1.226 | 0.924 | N/R | 0.830 | N/R |
| DeepAtom<sup>121</sup> | 1.232 | 0.904 | N/R | 0.831 | N/R |
| GraphscoreDTA<sup>37</sup> | 1.249 | 0.981 | N/R | 0.831 | N/R |
| EGNA<sup>43</sup> | 1.258 | 0.980 | N/R | 0.842 | N/R |
| PerSpect ML<sup>122</sup> | 1.265 | N/R | N/R | 0.840 | N/R |
| GlaNt<sup>40</sup> | 1.269 | 0.999 | N/R | 0.814 | N/R |
| KDEEP<sup>30</sup> | 1.27 | N/R | N/R | 0.82 | 0.82 |
| AGL-Score<sup>123</sup> | 1.272 | N/R | N/R | 0.833 | N/R |
| OnionNet<sup>25</sup> | 1.278 | 0.984 | N/R | 0.816 | N/R |
| PSH-GBT<sup>124</sup> | 1.280 | N/R | N/R | 0.835 | N/R |
| ELGN<sup>47</sup> | 1.285 | 1.013 | N/R | 0.810 | N/R |
| FAST<sup>35</sup> | 1.308 | 1.019 | 0.638 | 0.810 | 0.807 |
| BAPA<sup>125</sup> | 1.308 | 1.021 | N/R | 0.819 | 0.819 |
| SIGN<sup>36</sup> | 1.316 | 1.027 | N/R | 0.797 | N/R |
| TopologyNet<sup>23</sup> | 1.34 | N/R | N/R | 0.81 | N/R |
| DockingApp RF<sup>20</sup> | 1.35 | 1.09 | N/R | 0.83 | N/R |
| DeepDTAF<sup>126</sup> | 1.355 | 1.073 | N/R | 0.789 | N/R |
| DLSSAffinity<sup>127</sup> | 1.40 | N/R | N/R | 0.79 | N/R |
| DeepBindGCN<sup>49</sup> | 1.41 | N/R | N/R | 0.75 | N/R |
| Pafnucy<sup>31</sup> | 1.42 | 1.13 | N/R | 0.78 | N/R |
| Pair<sup>128</sup> | 1.44 | N/R | N/R | 0.75 | N/R |
| GraphBAR<sup>41</sup> | 1.542 | 1.241 | N/R | 0.726 | N/R |
| PointTransformer<sup>69</sup> | 1.58 | 1.29 | N/R | 0.753 | 0.751 |
| MGNN<sup>129</sup> | N/R | N/R | N/R | 0.85 | N/R |
| SE-OnionNet<sup>27</sup> | N/R | N/R | N/R | 0.83 | N/R |
| PLEC-NN<sup>130</sup> | N/R | N/R | N/R | 0.817 | N/R |

<sup>a</sup> The table reports Root Mean Square Error (RMSE), Mean Absolute Error (MAE), coefficient of determination ( $r^2$ ), Pearson correlation coefficient ( $r$ ), and Spearman rank correlation coefficient ( $\rho$ ) for each model.
<sup>b</sup> T-ALPHA<sup>†</sup> represents the performance of T-ALPHA evaluated on protein-ligand complex structures generated by Chai1 rather than the crystal structures.
<sup>c</sup> The best value for each metric is shown in bold.
<sup>d</sup> Models are sorted by RMSE value in ascending order.
<sup>e</sup> Error metrics (RMSE and MAE) are reported in units of $\text{p}K_i / \text{p}K_d$ .
<sup>f</sup> N/R indicates not reported in the literature.
<sup>g</sup> Values are reported with the number of significant figures provided in the original work.

In real-world drug discovery applications, experimental structures are often unavailable or incomplete. To evaluate the robustness of T-ALPHA in such scenarios, we assessed its performance on protein-ligand complex structures generated by Chai1 (T-ALPHA^†^). The results indicate that T-ALPHA^†^ maintains excellent performance (RMSE: 1.134, MAE: 0.893, *r*^2^: 0.721, Pearson *r*: 0.857, Spearman *ρ*: 0.843), outperforming all existing models in the literature that were evaluated using crystal structures (Figure S6 in the *Supporting Information*).

### 6.2 Assessing Generalizability Using LP-PDBbind and BDB2020+

To evaluate the ability of T-ALPHA to generalize to protein-ligand complexes significantly distinct from those in the training and validation distributions, we benchmarked its performance on two complementary test sets developed to assess generalizability of protein-ligand binding affinity scoring functions: LP-PDBbind, which was designed to evaluate internal generalizability by minimizing overlap between training, validation and test sets constructed from PDBbind data, and BDB2020+, which was curated to assess external generalizability to data obtained independently of PDBbind and collected after the training and validation data (More details in Section 2.1.1).

On the LP-PDBbind test set, T-ALPHA outperforms all previously evaluated models (Table S7 and Figure S8 in the *Supporting Information*).

On the BDB2020+ test set, T-ALPHA achieves an RMSE of 0.969, significantly outperforming all other models that have been evaluated (Table 4; Figure S9 in the *Supporting Information*). When using the Chai1-generated structures of the test set protein-ligand complexes rather than the crystal structures, the RMSE is also lower than that of any model previously evaluated on the crystal structures (Table 4; Figure S10 in the *Supporting Information*).

**Table 4.** Performance of T-ALPHA and Competing Models on the BDB2020+ Test Set.

| Model | RMSE | MAE | $r^2$ | Pearson $r$ | Spearman $\rho$ |
| --- | --- | --- | --- | --- | --- |
| T-ALPHA | 0.969 | 0.768 | 0.259 | **0.683** | 0.531 |
| T-ALPHA<sup>†</sup> | **0.939** | **0.740** | **0.310** | 0.681 | **0.544** |
| AutoDock Vina | 1.54 | N/R | N/R | 0.29 | N/R |
| IGN | 1.01 | N/R | N/R | 0.54 | N/R |
| RF-Score | 1.18 | N/R | N/R | 0.51 | N/R |
| DeepDTA | 1.26 | N/R | N/R | 0.26 | N/R |

<sup>a</sup> The table reports Root Mean Square Error (RMSE), Mean Absolute Error (MAE), coefficient of determination ( $r^2$ ), Pearson correlation coefficient ( $r$ ), and Spearman rank correlation coefficient ( $\rho$ ) for each model.
<sup>b</sup> T-ALPHA<sup>†</sup> represents the performance of T-ALPHA evaluated on protein-ligand complex structures generated by Chai1 rather than the crystal structures.
<sup>c</sup> The best value for each metric is shown in bold.
<sup>d</sup> Error metrics (RMSE and MAE) are reported in units of $pK_i / pK_d$ .
<sup>e</sup> N/R indicates not reported in the literature.

### 6.3 Protein-Specific Evaluations Using Mpro and EGFR Test Sets

In many real-world drug discovery scenarios, ranking compounds by binding affinity against a specific protein target is a critical step in prioritizing candidates for further investigation. In this context, a key metric is the Spearman rank correlation coefficient (*ρ*), which quantifies the monotonic relationship between the rank values of predicted and experimentally determined binding affinities.

To assess T-ALPHA’s effectiveness in protein-specific applications, we independently tested its performance for two highly relevant protein targets: SARS-CoV-2 main protease (Mpro) and epidermal growth factor receptor (EGFR). For the Mpro test set, T-ALPHA achieved a Spearman *ρ* of 0.737, outperforming all other models that have been evaluated (Table 5; Figure S11 in the *Supporting Information*). Even with the Chai1-generated structures (T-ALPHA^†^), the model maintained a Spearman *ρ* of 0.733 (Figure S12 in the *Supporting Information*).

**Table 5.**
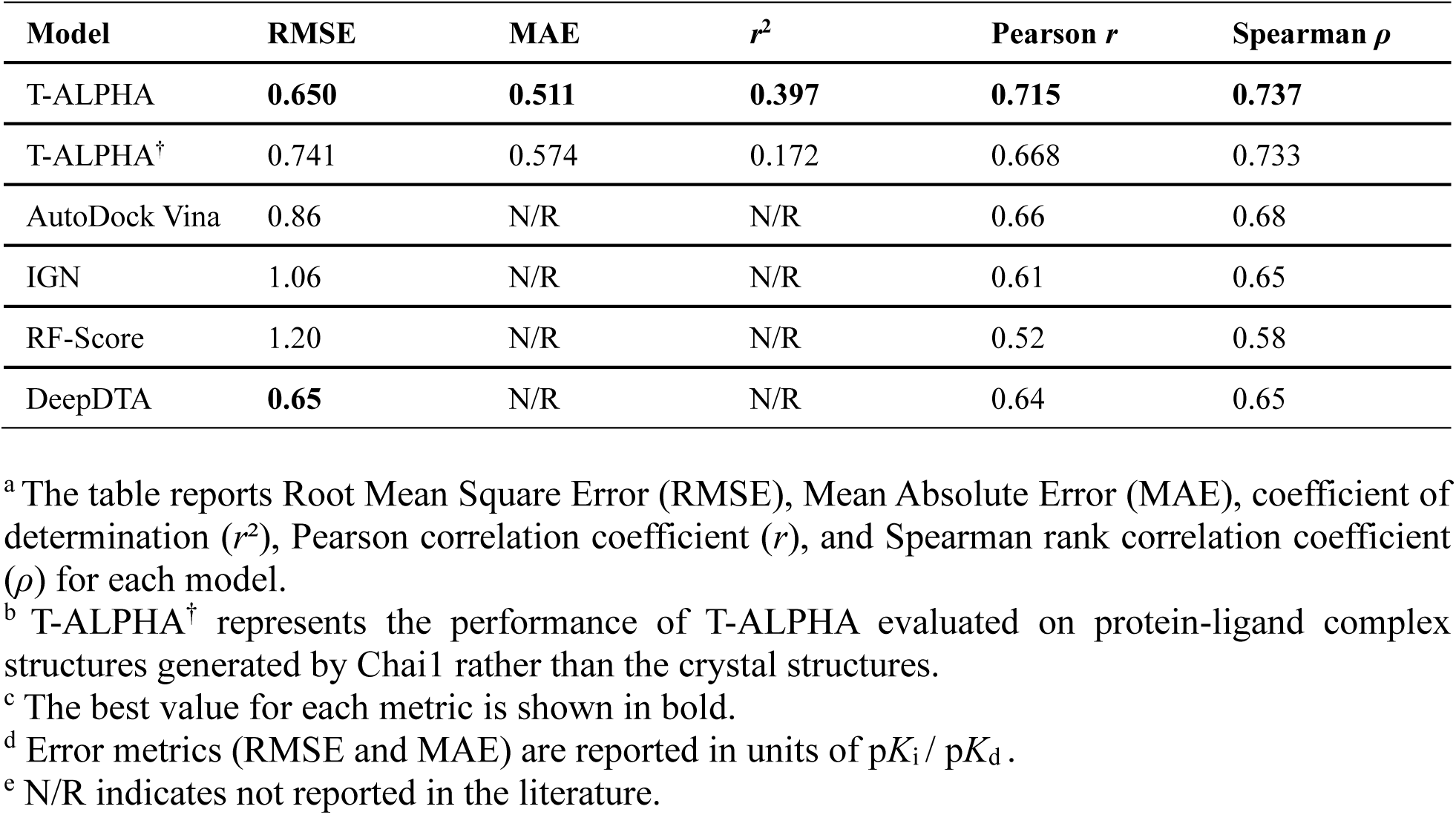
Performance of T-ALPHA and Competing Models on the Mpro Test Set.

| Model | RMSE | MAE | $r^2$ | Pearson $r$ | Spearman $\rho$ |
| --- | --- | --- | --- | --- | --- |
| T-ALPHA | **0.650** | **0.511** | **0.397** | **0.715** | **0.737** |
| T-ALPHA<sup>†</sup> | 0.741 | 0.574 | 0.172 | 0.668 | 0.733 |
| AutoDock Vina | 0.86 | N/R | N/R | 0.66 | 0.68 |
| IGN | 1.06 | N/R | N/R | 0.61 | 0.65 |
| RF-Score | 1.20 | N/R | N/R | 0.52 | 0.58 |
| DeepDTA | **0.65** | N/R | N/R | 0.64 | 0.65 |

<sup>a</sup> The table reports Root Mean Square Error (RMSE), Mean Absolute Error (MAE), coefficient of determination ( $r^2$ ), Pearson correlation coefficient ( $r$ ), and Spearman rank correlation coefficient ( $\rho$ ) for each model.
<sup>b</sup> T-ALPHA<sup>†</sup> represents the performance of T-ALPHA evaluated on protein-ligand complex structures generated by Chai1 rather than the crystal structures.
<sup>c</sup> The best value for each metric is shown in bold.
<sup>d</sup> Error metrics (RMSE and MAE) are reported in units of $pK_i / pK_d$ .
<sup>e</sup> N/R indicates not reported in the literature.

For the EGFR test set, T-ALPHA achieved a Spearman *ρ* of 0.791, significantly exceeding that of all other models evaluated (Table 6; Figures S13 and S14 in the *Supporting Information*).

**Table 6.** Performance of T-ALPHA and Competing Models on the EGFR Test Set.

| Model | RMSE | MAE | $r^2$ | Pearson $r$ | Spearman $\rho$ |
| --- | --- | --- | --- | --- | --- |
| T-ALPHA | **0.694** | **0.572** | **0.403** | **0.702** | **0.791** |
| T-ALPHA<sup>†</sup> | 0.842 | 0.670 | 0.093 | 0.593 | 0.665 |
| AutoDock Vina | 1.17 | N/R | N/R | 0.38 | 0.36 |
| IGN | 0.70 | N/R | N/R | 0.65 | 0.62 |
| RF-Score | 0.71 | N/R | N/R | 0.52 | 0.45 |
| DeepDTA | 0.77 | N/R | N/R | 0.44 | 0.43 |

<sup>a</sup> The table reports Root Mean Square Error (RMSE), Mean Absolute Error (MAE), coefficient of determination ( $r^2$ ), Pearson correlation coefficient ( $r$ ), and Spearman rank correlation coefficient ( $\rho$ ) for each model.
<sup>b</sup> T-ALPHA<sup>†</sup> represents the performance of T-ALPHA evaluated on protein-ligand complex structures generated by Chai1 rather than the crystal structures.
<sup>c</sup> The best value for each metric is shown in bold.
<sup>d</sup> Error metrics (RMSE and MAE) are reported in units of $\text{p}K_i / \text{p}K_d$ .
<sup>e</sup> N/R indicates not reported in the literature.

### 6.4 Enhancing Target-Specific Performance with Self-Learning

We assessed the effectiveness of the proposed self-learning method by comparing T-ALPHA’s performance on the Mpro and EGFR test sets before and after its application. For the Mpro test set, the newly trained model and fine-tuned model demonstrate significant improvements in Spearman *ρ* compared to the baseline, with increases of 9.91% and 5.43%, respectively (Table S15 in the *Supporting Information*). Importantly, the control experiments exhibit minimal change compared to the baseline, confirming that the observed improvements are attributable to the self-learning method rather than other factors. Similar trends are observed for application of the method to EGFR, with increases in Spearman *ρ* of 3.41% for the newly trained model and 1.14% for the fine-tuned model (Table S16 in the *Supporting Information*). These results highlight the effectiveness of the proposed method in enhancing protein-specific ranking of compounds by binding affinity, although the magnitude of improvement varies depending on the specified target.

An interesting and somewhat unintuitive observation is that although Spearman *ρ* and Pearson *r* increase for the applications of the method to both Mpro and EGFR, the RMSE and MAE values also increase. This discrepancy arises due to the systematic biases inherent in the pseudo-labels generated by the model. As absolute metrics, RMSE and MAE are sensitive to these biases, whereas correlation metrics like Spearman *ρ* and Pearson *r*, which measure relative relationships, are largely unaffected.

### 6.5 Architecture Component Contributions to Performance

To validate the design choices underpinning the T-ALPHA architecture, we conducted an ablation study where we systematically removed individual components and characterized the relative contribution of each component to the overall performance on the CASF 2016 benchmark (Table 7).

**Table 7.**
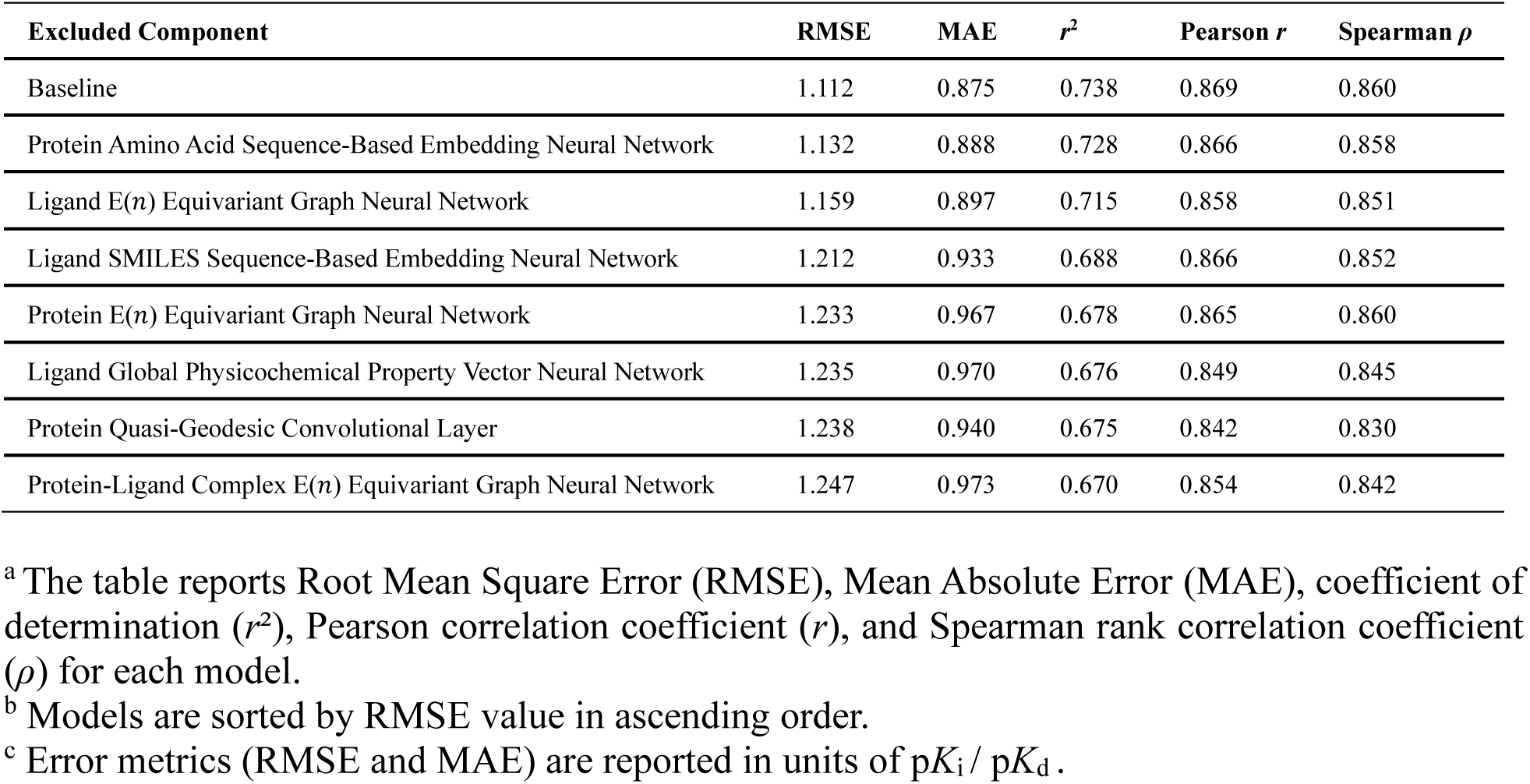
Impact of Excluded Components on T-ALPHA Performance on the CASF 2016 Benchmark.

Removing the protein-ligand complex E(*n*) EGNN led to the most significant decline in performance among all components, with an RMSE of 1.247 (Table 7). This component explicitly models the interactions between the protein and ligand atoms, and the performance decline due to its exclusion demonstrates that capturing these intermolecular relationships is critical for accurately predicting binding affinity.

Excluding the protein quasi-geodesic convolutional layer resulted in an RMSE of 1.238 (Table 7), underscoring the importance of capturing the topography and curvature of the protein binding pocket. This layer provides critical insights into potential interaction hotspots and steric compatibility, which inform the other components of the protein channel through cross-attention mechanisms. Removing the ligand physicochemical property-based embedding neural network led to an RMSE of 1.235, demonstrating that molecular-level properties of the ligand are informative of its binding behavior. Moreover, in the baseline model, the output of this component informs the other components of the ligand channel through cross-attention mechanisms.

The other components, including the protein and ligand E(*n*) EGNNs, as well as the protein and ligand sequence-based neural networks, each contributes meaningfully to the model’s performance (Table 7). The observed declines in performance due to their removals highlight the importance of connectivity-based structural features of both the protein pocket and the ligand, patterns describing chemical and functional relationships of the ligand, and global evolutionary and functional information about the entire protein for predicting binding affinity.

The ablation study reveals that all components contribute to the overall performance of T-ALPHA, with certain modules having a more substantial impact when excluded. These results confirm that the multimodal feature representations and hierarchical transformer architecture of T-ALPHA are critical to its state-of-the-art performance. Each component captures distinct aspects of protein-ligand interactions, and their integration enables the model to account for the diverse factors that determine binding affinity.

## 7 Discussion

In this work, we introduced T-ALPHA, a novel deep learning model designed to predict protein-ligand binding affinity by integrating multimodal feature representations through a hierarchical transformer framework. Our extensive evaluations demonstrate that T-ALPHA achieves state-of-the-art performance across multiple benchmarks, highlighting its applicability to early-stage drug discovery workflows.

On the widely recognized CASF 2016 benchmark, T-ALPHA outperforms all existing models reported in the literature, underscoring the effectiveness of our architectural approach in capturing the characteristics and interactions that determine binding affinity. Notably, even when using predicted protein-ligand complex structures rather than crystal structures, the model maintains superior performance to existing models. This robustness is particularly important in real-world drug discovery projects, where experimentally determined structures are often unavailable or incomplete.

In addition, T-ALPHA demonstrates effective generalizability via the LP-PDBbind and BDB2020+ benchmarks, which were specifically designed to evaluate performance on protein-ligand complexes outside of the training distribution, outperforming all models that were previously evaluated.

T-ALPHA also achieves state-of-the-art performance on protein-specific test sets corresponding to SARS-CoV-2 main protease (Mpro) and the epidermal growth factor receptor (EGFR), effectively ranking compounds by binding affinity to each of the respective targets—a key requirement in prioritization and lead optimization in drug discovery pipelines. Moreover, we proposed an uncertainty-aware self-learning method for protein-specific alignment that does not require additional experimental data, and demonstrated that it enhances the ability of T-ALPHA to rank compounds by binding affinity to both of the targets.

The ablation study performed revealed that each component of the architecture contributes to the overall accuracy of the model, validating the architectural design of T-ALPHA and highlighting the importance of multimodal feature integration for state-of-the-art performance.

## 8 Outlook and Future Directions

While T-ALPHA advances the field of protein-ligand binding affinity prediction, several challenges and opportunities remain. One of the foremost challenges is the availability of high-quality, standardized datasets that comprehensively cover the vast chemical and biological space. Current datasets suffer from inconsistencies in experimental techniques used to obtain binding affinity measurements, and limited coverage of diverse chemical structures and protein targets. Improving data quality through standardized experimental protocols, as well as expanding datasets to include a wider range of chemical entities and protein families, will significantly improve the capabilities of these models.

Despite progress, models often struggle to generalize to chemical and biological spaces beyond those represented in the training data. Developing methods to enhance generalizability, including leveraging transfer learning, zero-shot learning, and incorporating domain knowledge to guide predictions, is critical for advancing the field.

Reproducibility remains an area of ongoing improvement in the field, as many published models lack accessible or functional code, making it difficult to validate or build upon prior work. To address this gap, researchers should release fully functional code with clear documentation alongside their publications. To promote transparency and reproducibility, we have made all of our code and trained models openly available at https://github.com/gregory-kyro/T-ALPHA, enabling researchers to run T-ALPHA and reproduce all of the results presented in this paper.

## Software and Implementation

The implementation of T-ALPHA was conducted using PyTorch (v2.4.1+cu121)^131^ and PyTorch Geometric (v2.6.0).^132^ Training was performed with PyTorch Lightning.^115^ Model training employed features such as *ModelCheckpoint* for saving the best-performing models and *CSVLogger* for logging training metrics. Data handling leveraged PyTorch Geometric’s *DataListLoader* to batch and process graph-based datasets efficiently.

The E(*n*) EGNNs utilized PyTorch’s core modules (*torch* and *torch.nn*) to define custom neural network layers. The dMaSIF-based component combined PyTorch, PyTorch Geometric, and KeOps.^133^ The meta transformer architecture was implemented using PyTorch’s *TransformerEncoder* and *TransformerDecoder* modules, with graph-level pooling operations handled by PyTorch Geometric’s *global_mean_pool*. Training optimization incorporated learning rate schedulers including a combination of *LinearLR* for warm-up and *CosineAnnealingLR* for gradual learning rate decay. Metrics such as Pearson correlation coefficient were computed using SciPy^134^ for model evaluation.

Chai1 was run with three trunk cycles and 200 diffusion steps. The predicted structure with the best score was selected for downstream processing.

## Supporting Information

The Supporting Information contains details regarding the training and validation learning curves, custom loss function, CASF 2016 benchmark, LP-PDBbind benchmark, BDB2020+ benchmark, protein-specific Mpro and EGFR benchmarks, and the proposed uncertainty-aware self-learning method.

## Data and Software Availability

All of our code and trained models are openly available at https://github.com/gregory-kyro/T-ALPHA, enabling researchers to run T-ALPHA and reproduce all of the results presented in this paper.

## Acknowledgments

We acknowledge financial support from the National Science Foundation Graduate Research Fellowship under Grant DGE-2139841 [GWK], from the National Science Foundation Engines Development Award: Advancing Quantum Technologies (CT) under Award Number 2302908 [VSB], and from the CCI Phase I: National Science Foundation Center for Quantum Dynamics on Modular Quantum Devices (CQD-MQD) under Award Number 2124511 [VSB]. Additionally, we acknowledge high-performance computer time from the National Energy Research Scientific Computing Center and from the Yale University Faculty of Arts and Sciences High Performance Computing Center.

## Author Contributions

GWK, AMS, YS, CX, VSB conceived the idea; GWK, AMS, YS, CX designed research; GWK, AMS, YS, CX developed software; GWK, AMS, YS, CX performed research; GWK, AMS, YS, CX, VSB analyzed data; GWK, AMS, YS, CX wrote the paper; VSB provided feedback on the paper. All authors have given approval to the final version of the manuscript.

## Abbreviations

ML: machine learning
CNNs: convolutional neural networks
GNNs: graph neural networks
EGNN: equivariant graph neural network
Mpro: SARS-CoV-2 main protease
EGFR: epidermal growth factor receptor
NMR: nuclear magnetic resonance
*K*_i_: inhibition constant
*K*_d_: dissociation constant
IC_50_: half-maximal inhibitory concentration
CASF: Comparative Assessment of Scoring Functions
LP-PDBbind: Leak Proof PDBbind
MLP: multilayer perceptron
GELU: Gaussian Error Linear Unit
FSDP: fully sharded data parallel
ECFP4: extended connectivity fingerprint with a diameter of 4
RMSE: root mean square error
MAE: mean absolute error
*r*: Pearson correlation coefficient
*r*^2^: coefficient of determination
*ρ*: Spearman rank correlation coefficient.

## Supporting Information

**Figure S1.**
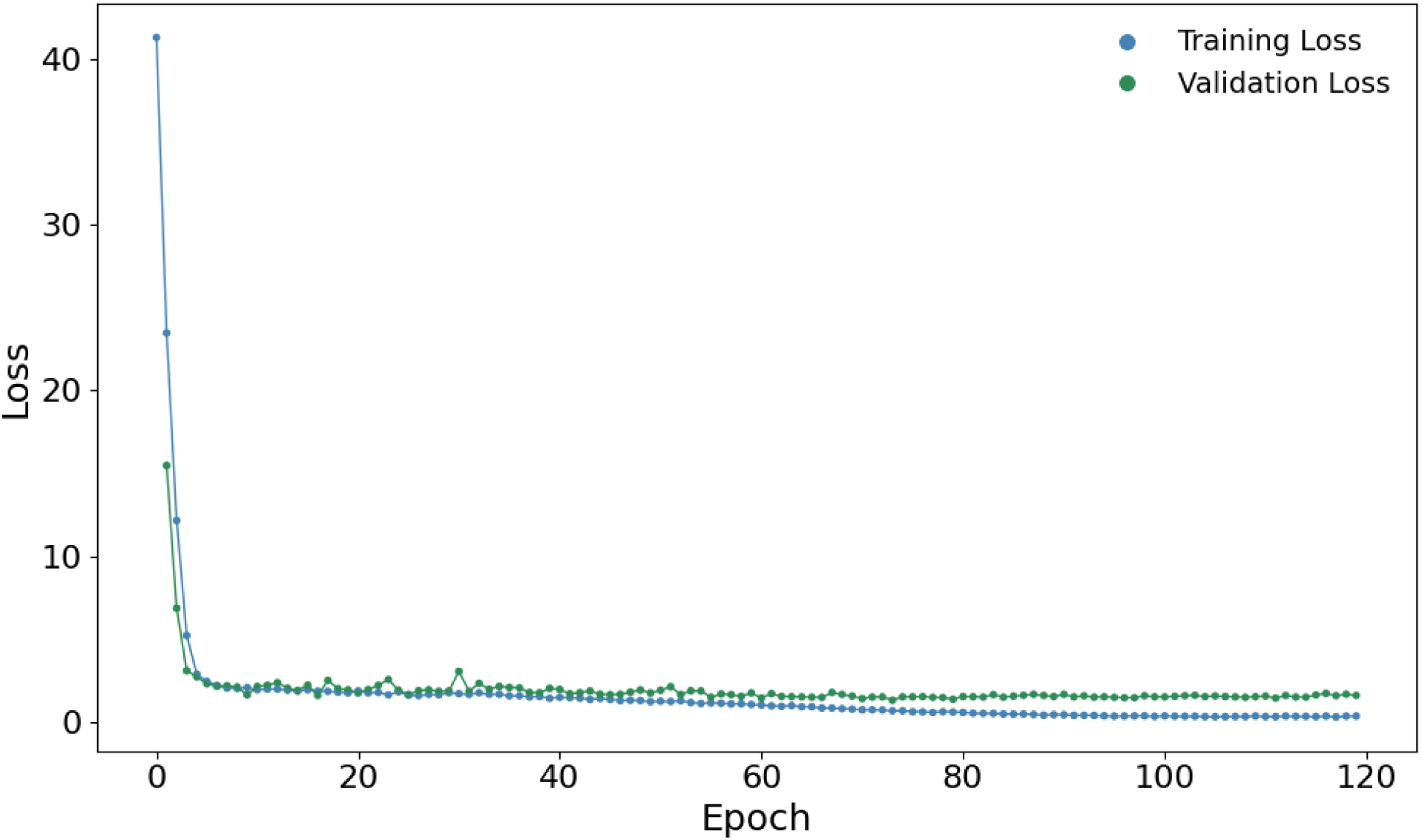
Training and validation loss curves for T-ALPHA corresponding to testing on the CASF 2016 test set. Loss is calculated using the custom loss function and plotted as a function of epoch number.

**Figure S2.**
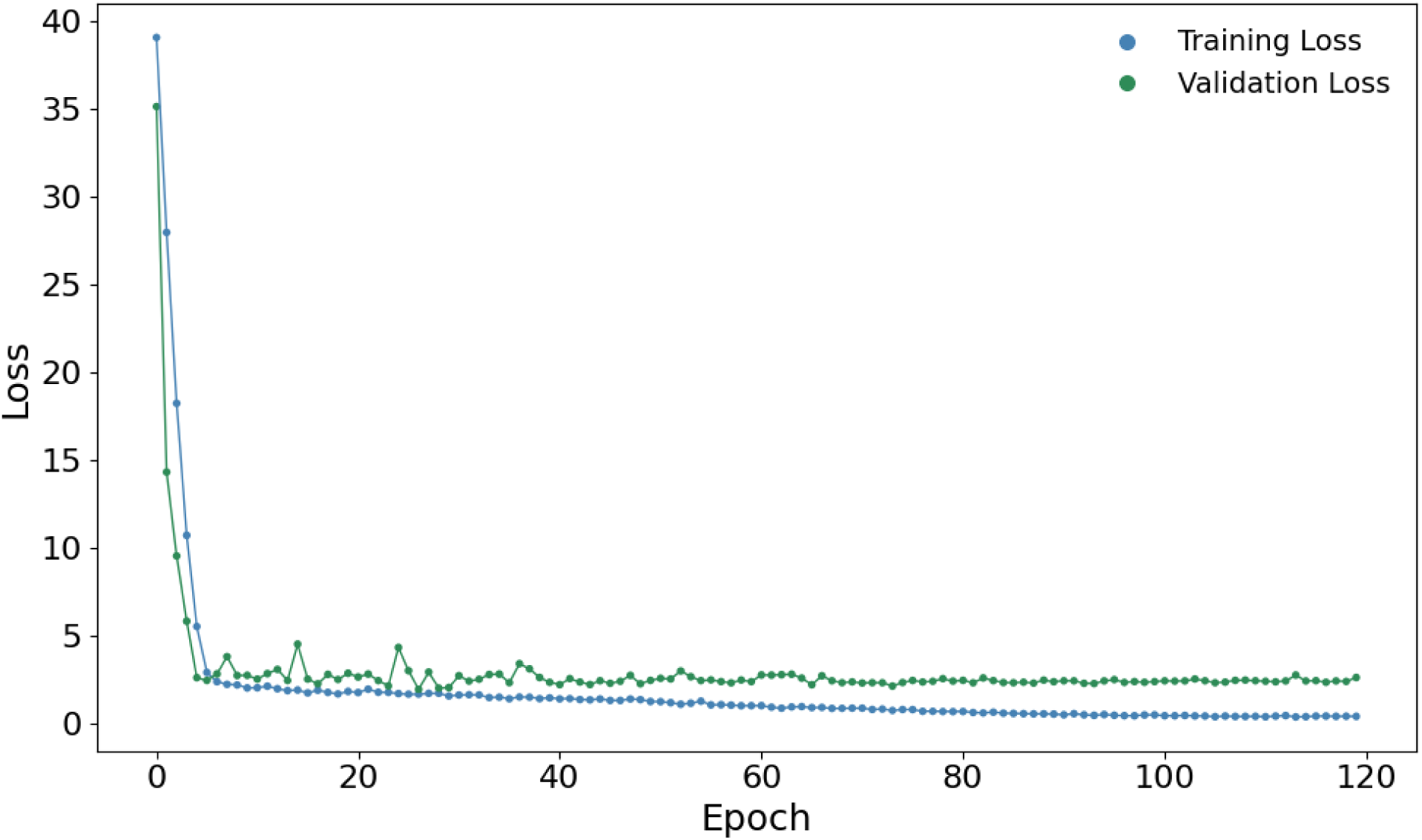
Training and validation loss curves for T-ALPHA corresponding to testing on the LP-PDBbind test set. Loss is calculated using the custom loss function and plotted as a function of epoch number.

**Figure S3.**
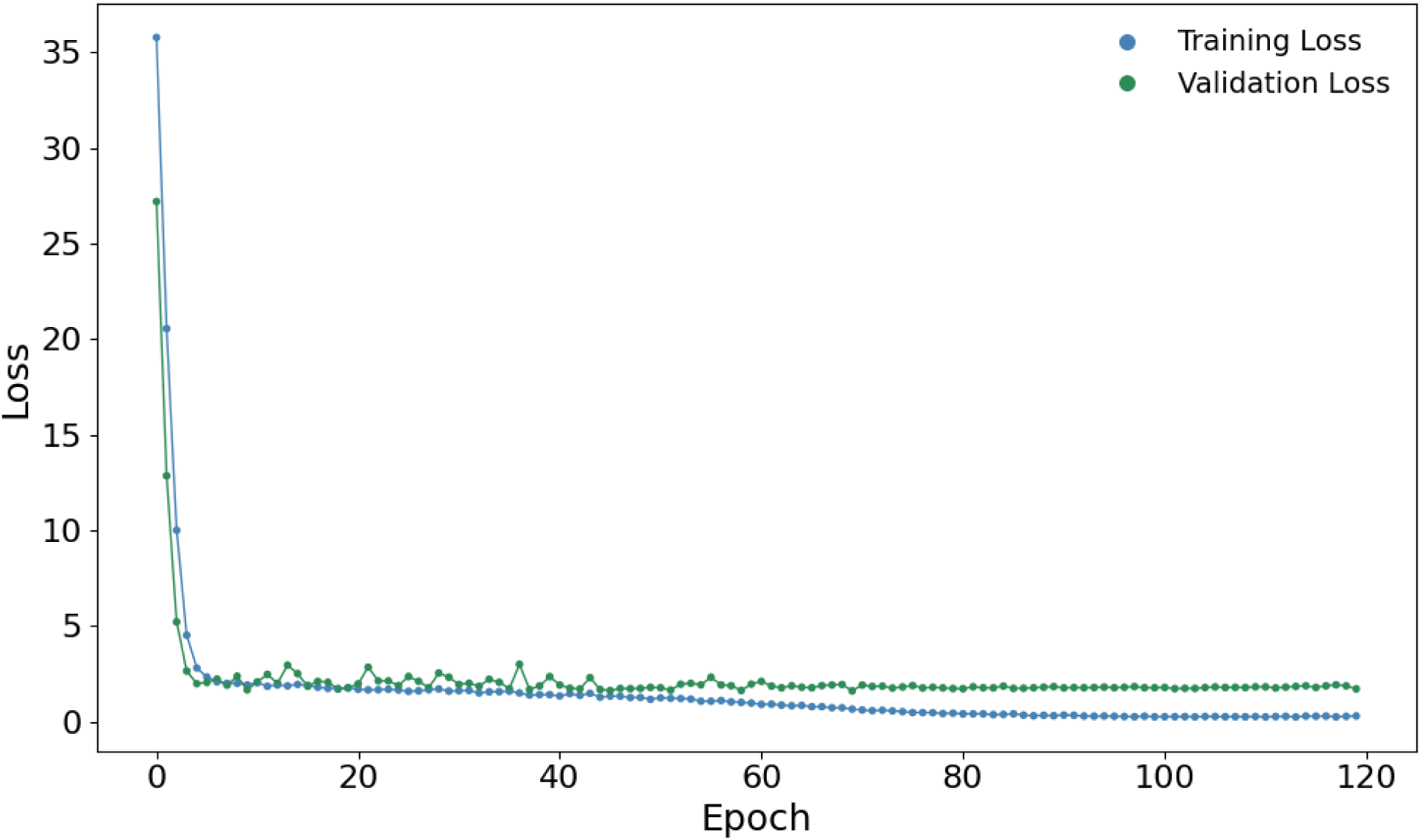
Training and validation loss curves for T-ALPHA corresponding to testing on the BDB2020+ test set, as well as protein-specific test sets for SARS-CoV-2 main protease (Mpro) and epidermal growth factor receptor (EGFR). Loss is calculated using the custom loss function and plotted as a function of epoch number.

**Figure S4.**
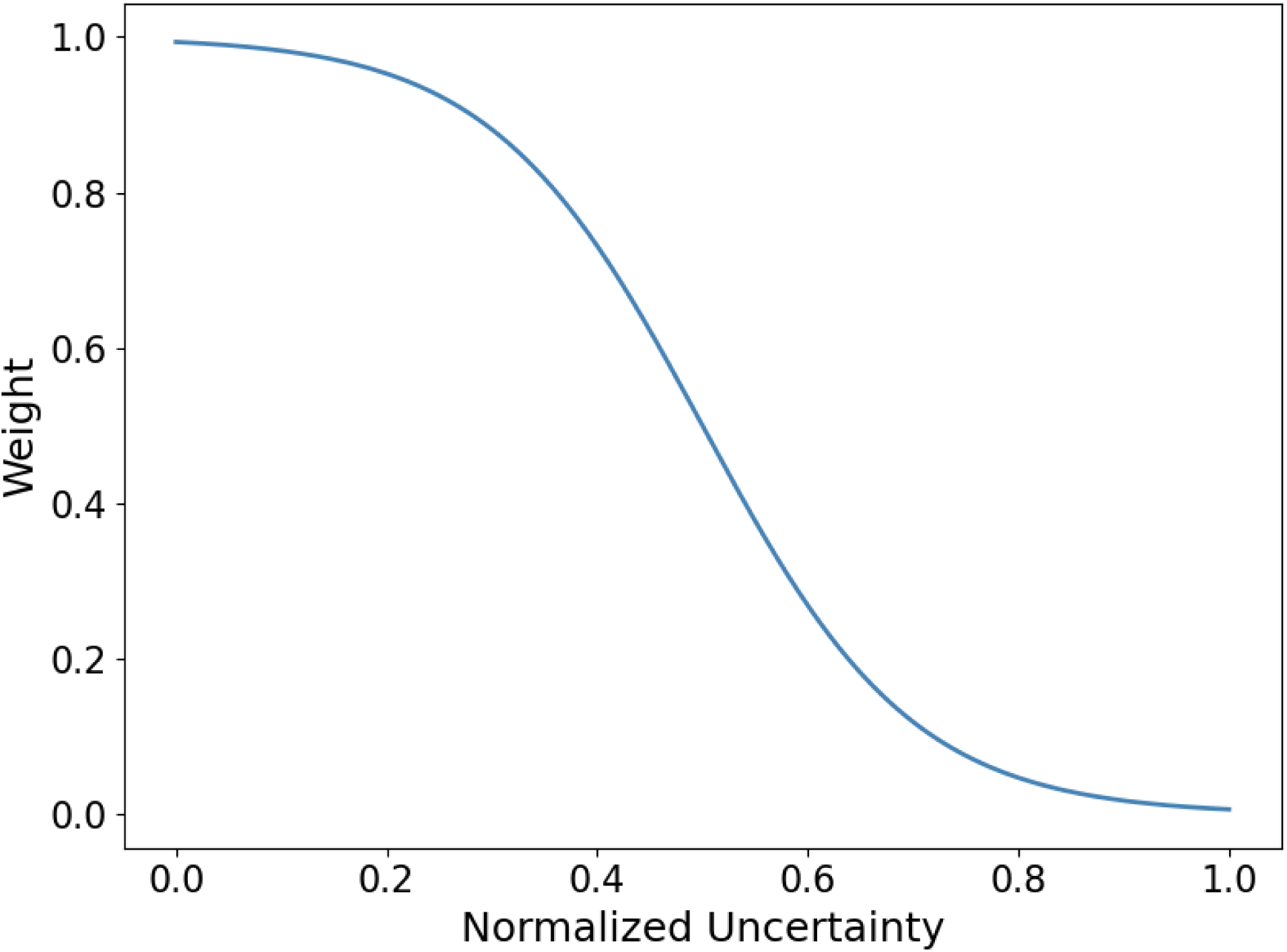
Weighting function for uncertainty-aware training. The figure shows the transformation applied to normalized uncertainty values to calculate weights for the loss function during parameter optimization. The function is defined as 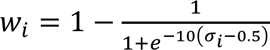, where *σ*_*i*_ is the normalized uncertainty. The transformation maps normalized uncertainty values to weights in the range (0, 1), smoothly decreasing the weight as uncertainty increases.

**Figure S5.**
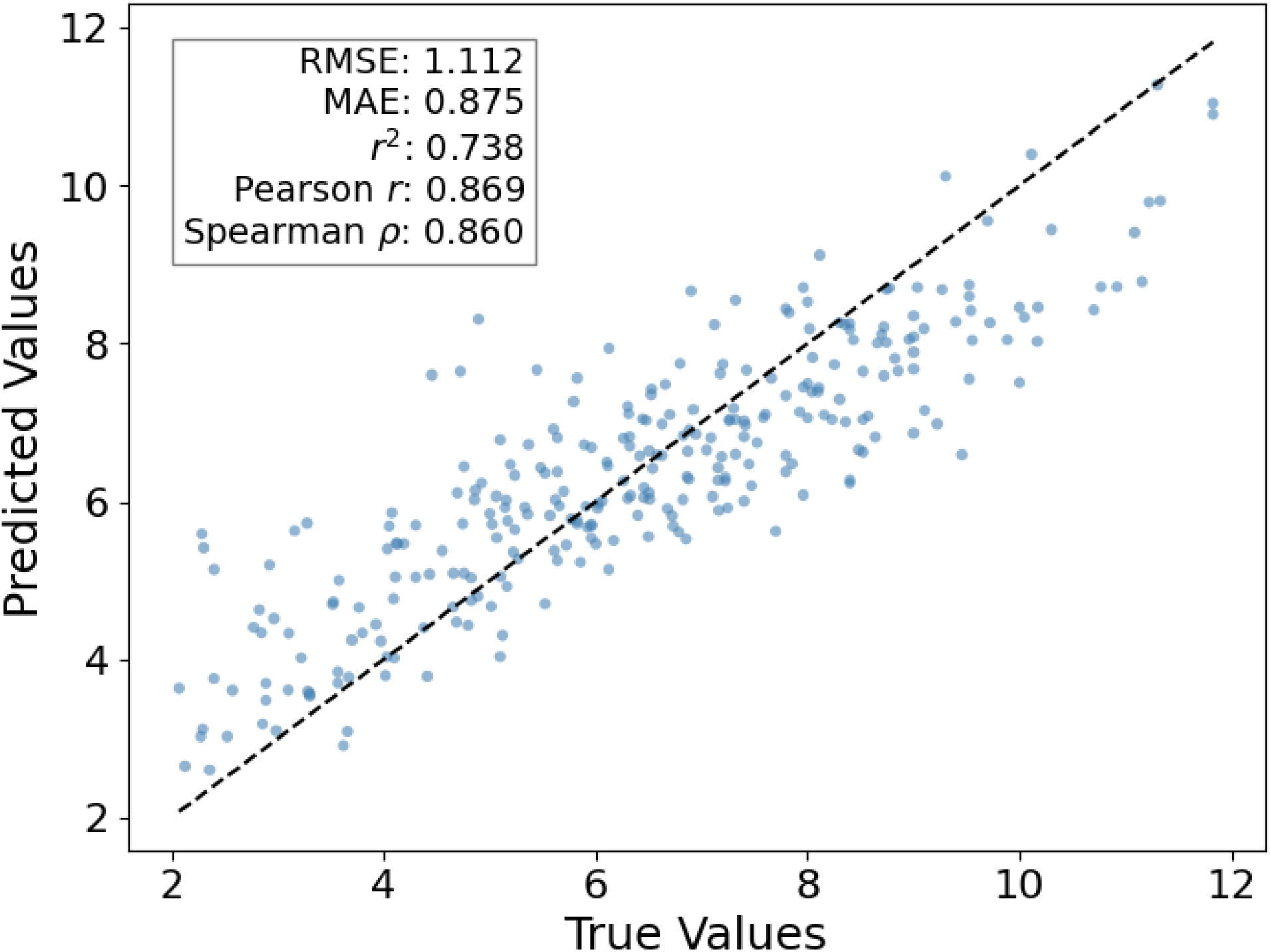
Predicted versus true binding affinities for T-ALPHA on the CASF 2016 test set using crystal structures of protein-ligand complexes. Performance metrics include Root Mean Square Error (RMSE), Mean Absolute Error (MAE), coefficient of determination (*r*²). Pearson correlation coefficient (*r*), and Spearman rank correlation coefficient (*ρ*).

**Figure S6.**
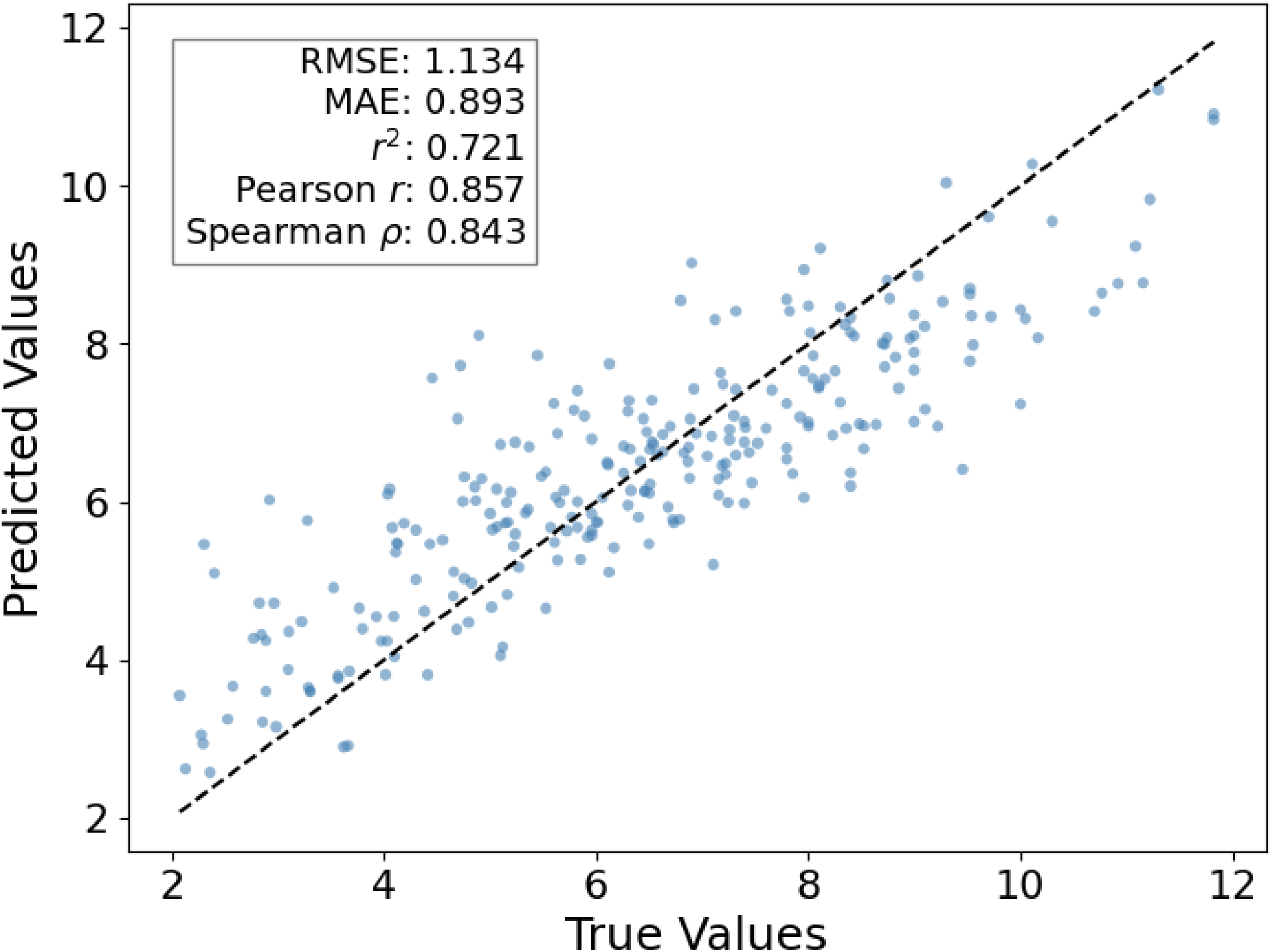
Predicted versus true binding affinities for T-ALPHA^†^ on the CASF 2016 test set using Chai1-generated protein-ligand complex structures. Performance metrics include Root Mean Square Error (RMSE), Mean Absolute Error (MAE), coefficient of determination (*r*²). Pearson correlation coefficient (*r*), and Spearman rank correlation coefficient (*ρ*).

**Table S7.**
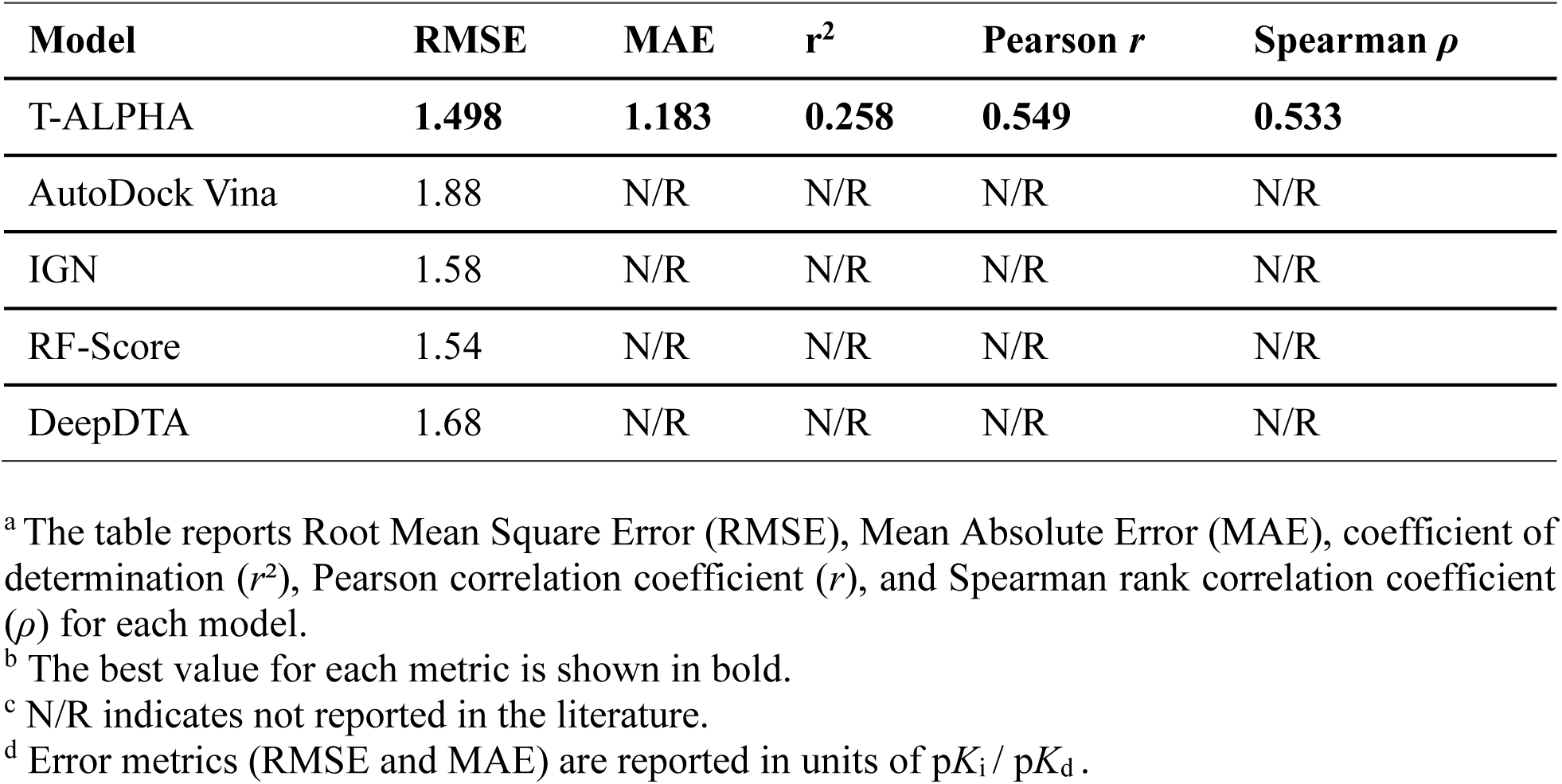
Performance of T-ALPHA and models reported in the literature on the LP-PDBbind test set.

**Figure S8.**
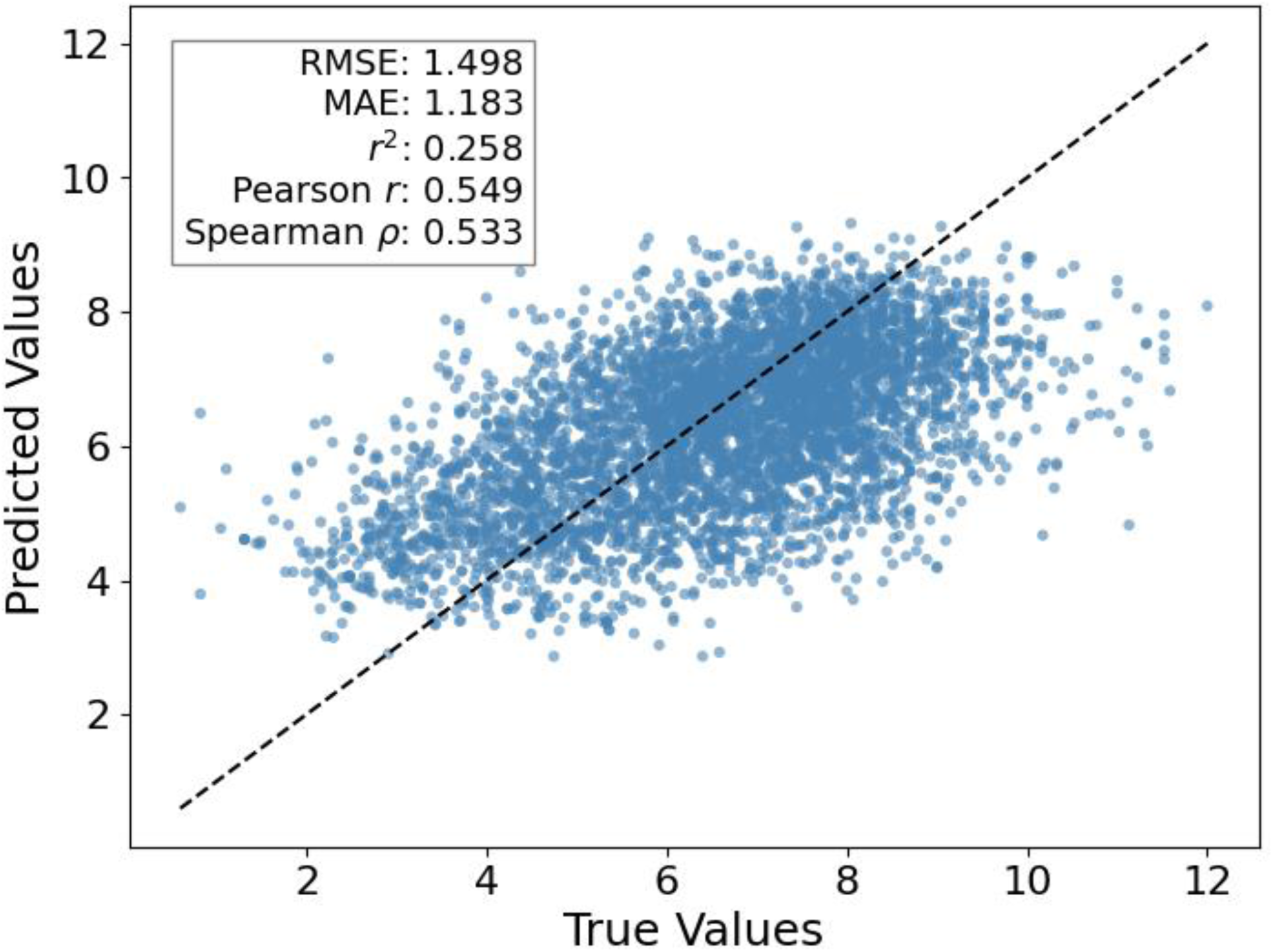
Predicted versus true binding affinities for T-ALPHA on the LP-PDBbind test set. Performance metrics include Root Mean Square Error (RMSE), Mean Absolute Error (MAE), coefficient of determination (*r*²). Pearson correlation coefficient (*r*), and Spearman rank correlation coefficient (*ρ*).

**Figure S9.**
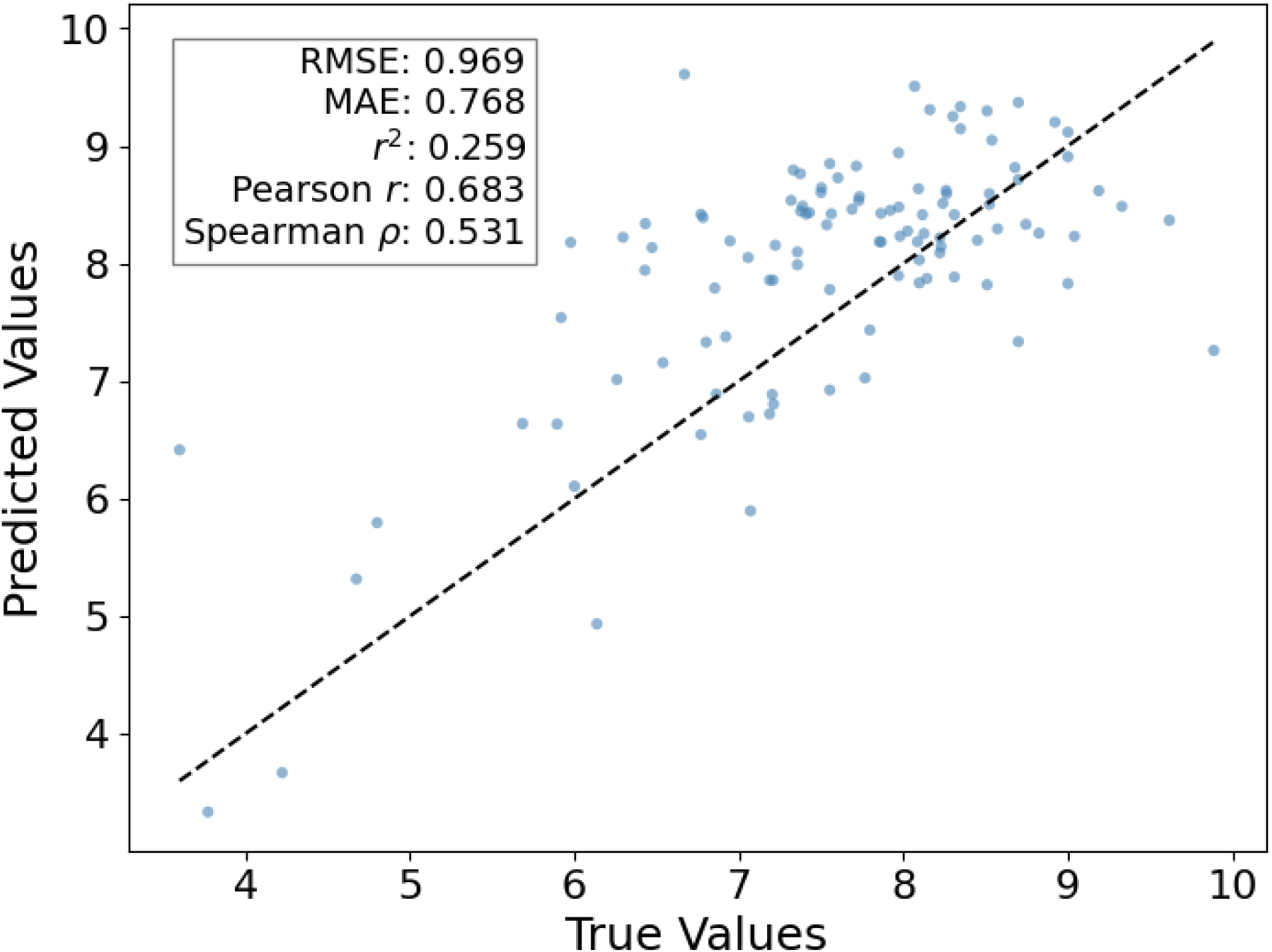
Predicted versus true binding affinities for T-ALPHA on the BDB2020+ test set. Performance metrics include Root Mean Square Error (RMSE), Mean Absolute Error (MAE), coefficient of determination (*r*²). Pearson correlation coefficient (*r*), and Spearman rank correlation coefficient (*ρ*).

**Figure S10.**
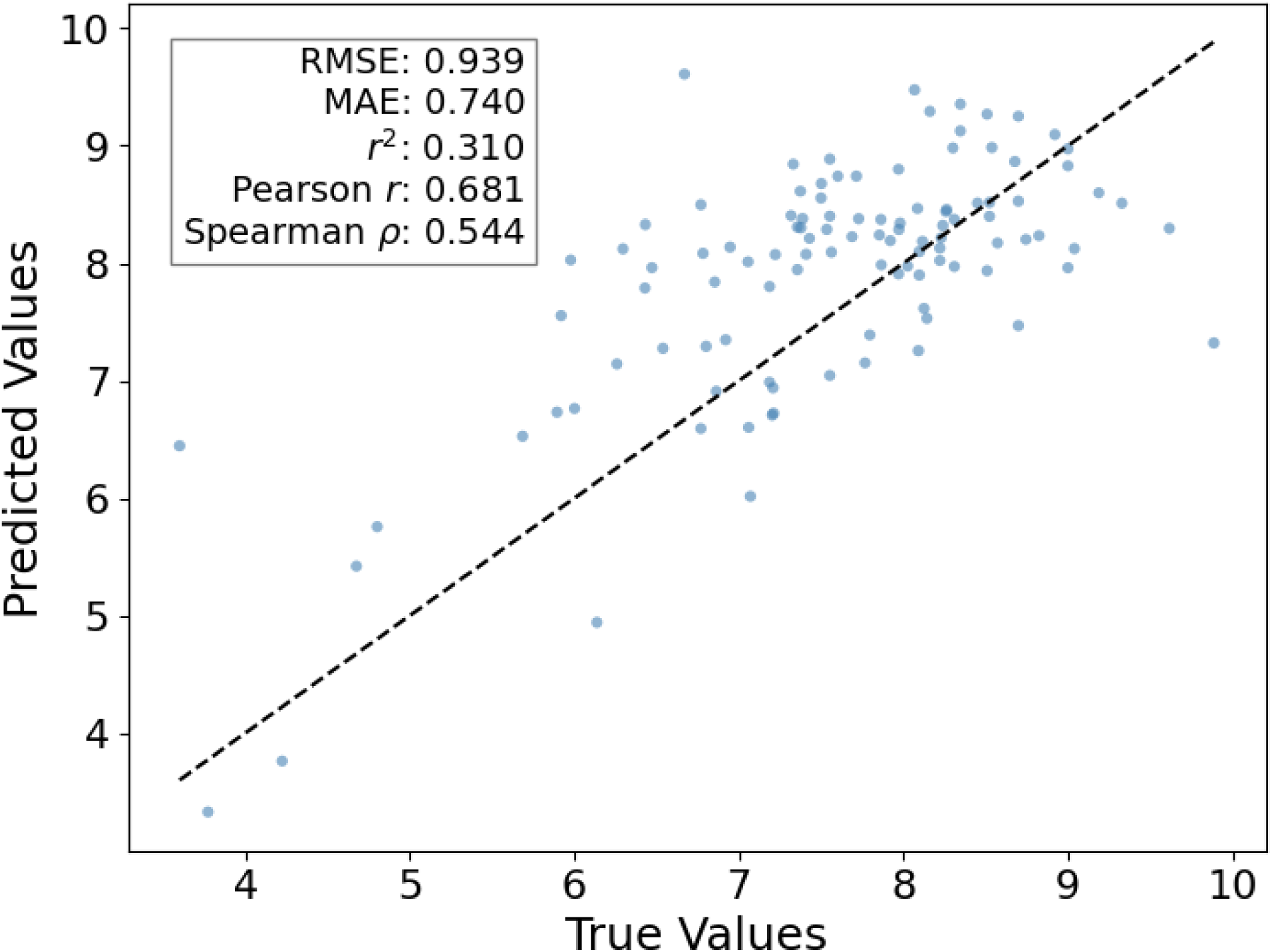
Predicted versus true binding affinities for T-ALPHA^†^ on the Chai1-generated protein-ligand complex structures of the BDB2020+ test set. Performance metrics include Root Mean Square Error (RMSE), Mean Absolute Error (MAE), coefficient of determination (*r*²). Pearson correlation coefficient (*r*), and Spearman rank correlation coefficient (*ρ*).

**Figure S11.**
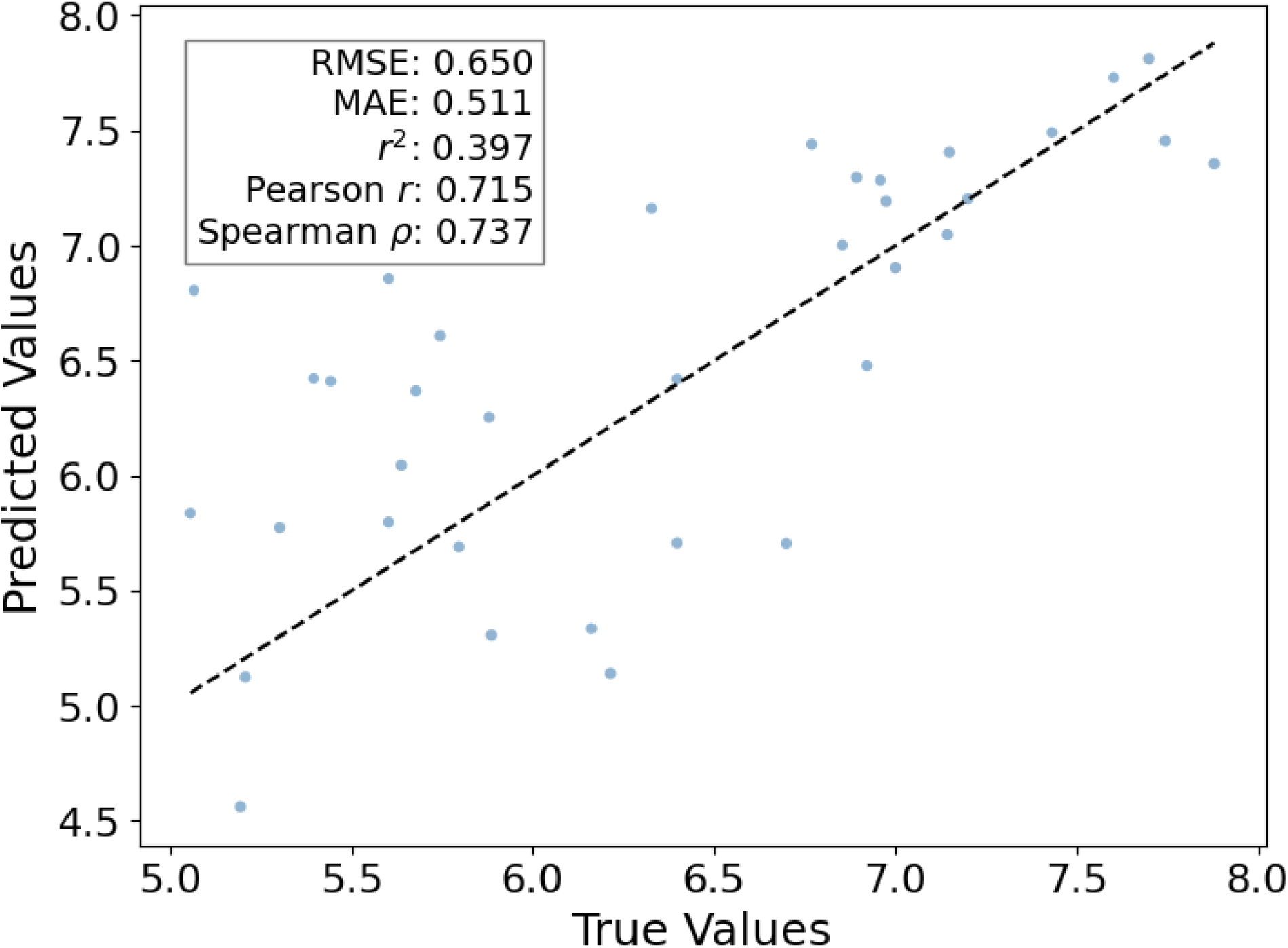
Predicted versus true binding affinities for T-ALPHA on the Mpro test set. Performance metrics include Root Mean Square Error (RMSE), Mean Absolute Error (MAE), coefficient of determination (*r*²). Pearson correlation coefficient (*r*), and Spearman rank correlation coefficient (*ρ*).

**Figure S12.**
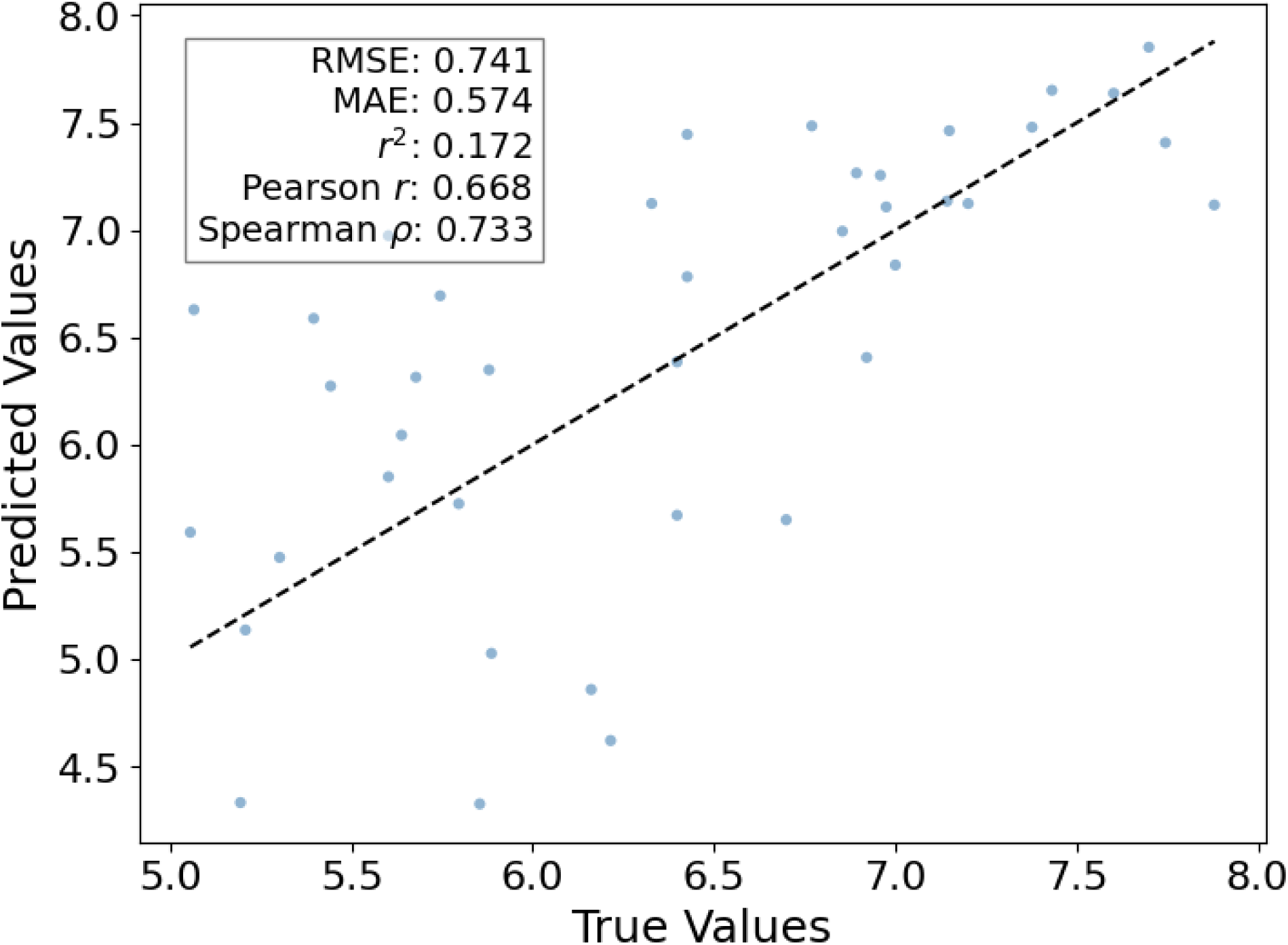
Predicted versus true binding affinities for T-ALPHA^†^ on the Chai1-generated protein-ligand complex structures of the Mpro test set. Performance metrics include Root Mean Square Error (RMSE), Mean Absolute Error (MAE), coefficient of determination (*r*²). Pearson correlation coefficient (*r*), and Spearman rank correlation coefficient (*ρ*).

**Figure S13.**
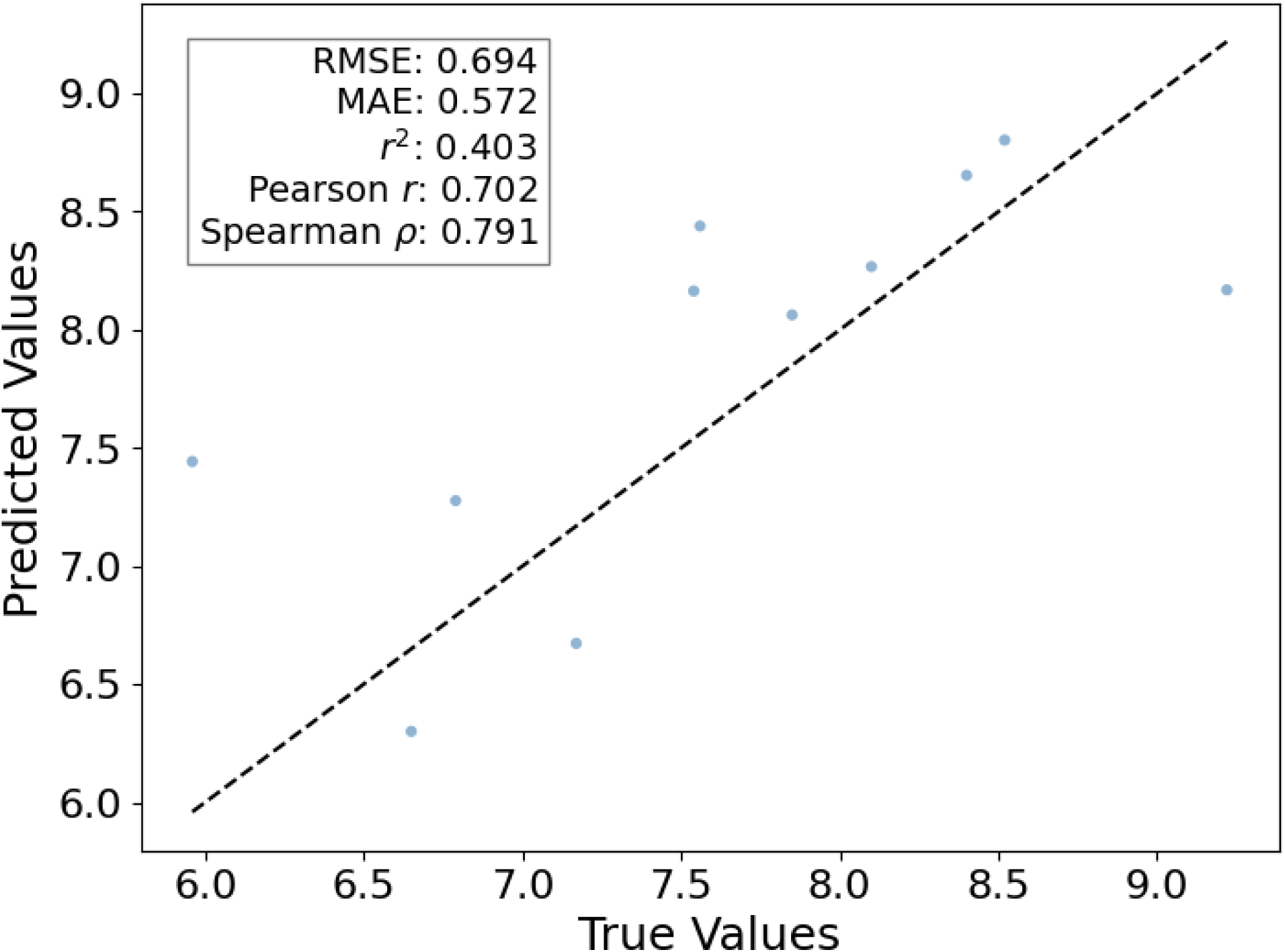
Predicted versus true binding affinities for T-ALPHA on the EGFR test set. Performance metrics include Root Mean Square Error (RMSE), Mean Absolute Error (MAE), coefficient of determination (*r*²). Pearson correlation coefficient (*r*), and Spearman rank correlation coefficient (*ρ*).

**Figure S14.**
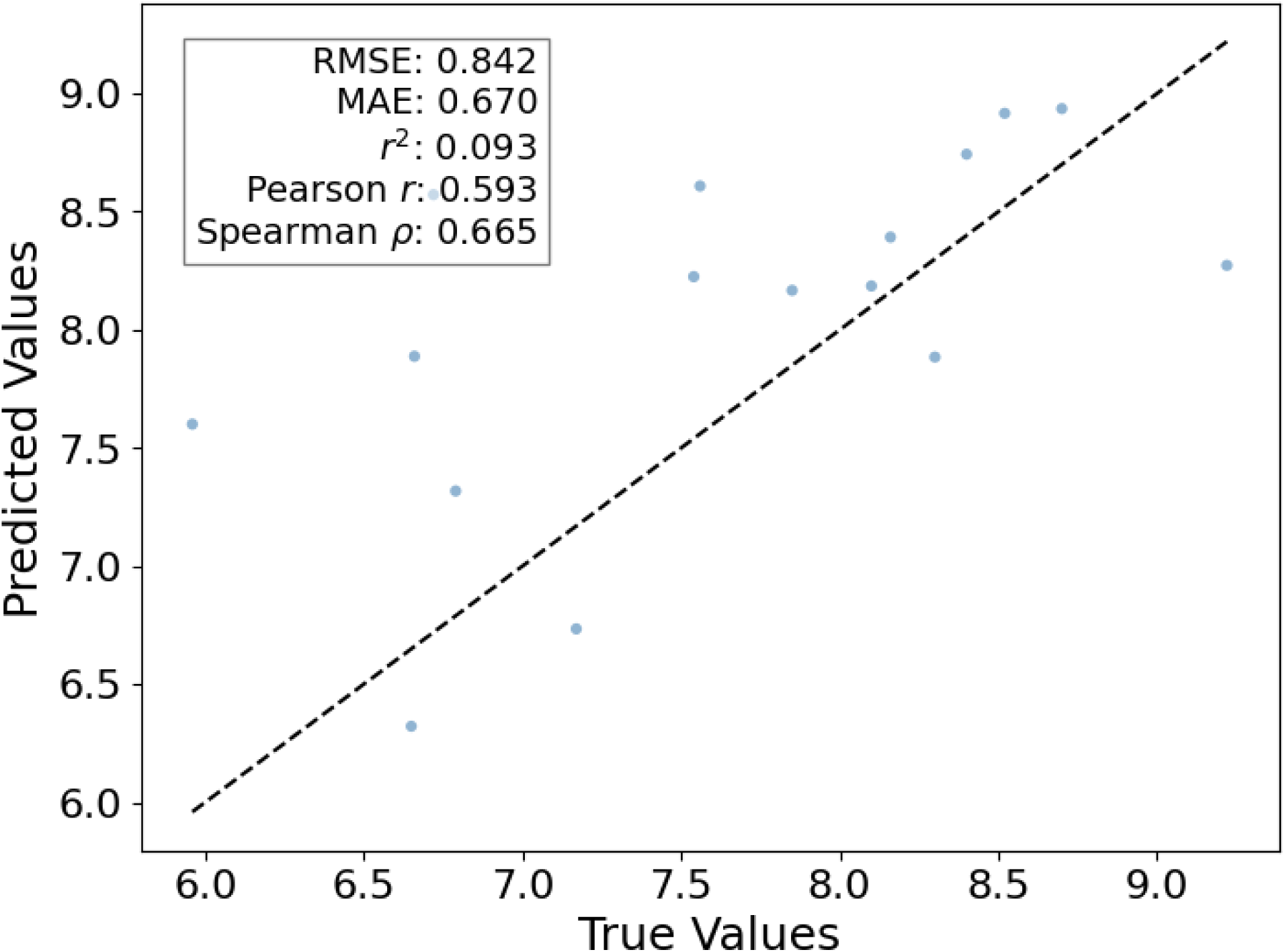
Predicted versus true binding affinities for T-ALPHA^†^ on the Chai1-generated protein-ligand complex structures of the EGFR test set. Performance metrics include Root Mean Square Error (RMSE), Mean Absolute Error (MAE), coefficient of determination (*r*²). Pearson correlation coefficient (*r*), and Spearman rank correlation coefficient (*ρ*).

**Table S15.** Improvements in Spearman’s rank correlation coefficient (*ρ*) for Mpro using the proposed self-learning method. Performance metrics are shown for comparing the baseline, new model, and fine-tuned model for SARS-CoV-2 main protease (Mpro). Spearman *ρ* improvements of 9.91% for the new model and 5.43% for the fine-tuned model highlight the effectiveness of the self-learning method. Results are reported separately for crystal structures (first value) and Chai1-generated structures (values in parentheses).

| **Model** | **RMSE** | **MAE** | **Pearson $r$** | **Spearman $\rho$** | **% $\uparrow \Delta$ Spearman $\rho$** |
| --- | --- | --- | --- | --- | --- |
| Baseline | 0.650<br>(0.741) | 0.511<br>(0.574) | 0.715<br>(0.668) | 0.737<br>(0.733) | 0.00% (0.00%) |
| New Model | 1.397<br>(1.468) | 1.259<br>(1.324) | 0.790<br>(0.776) | 0.810<br>(0.803) | 9.91% (9.55%) |
| Control (New Model) | 1.374<br>(1.387) | 1.210<br>(1.200) | 0.741<br>(0.700) | 0.737<br>(0.702) | 0.00% (-4.22%) |
| Fine Tune | 0.615<br>(0.651) | 0.529<br>(0.549) | 0.752<br>(0.718) | 0.777<br>(0.762) | 5.43% (3.96%) |
| Control (Fine Tune) | 0.915<br>(0.923) | 0.779<br>(0.776) | 0.764<br>(0.750) | 0.735<br>(0.732) | -0.27% (-0.14%) |

<sup>a</sup> The table reports Root Mean Square Error (RMSE), Mean Absolute Error (MAE), Pearson correlation coefficient ( $r$ ), Spearman rank correlation coefficient ( $\rho$ ), and the percentage increase in Spearman $\rho$ compared to the baseline (% $\uparrow \Delta$ Spearman $\rho$ ).
<sup>b</sup> Error metrics (RMSE and MAE) are reported in units of $pK_i / pK_d$ .

**Table S16.** Improvements in Spearman’s rank correlation coefficient (*ρ*) for EGFR using the proposed self-learning method. Performance metrics are shown for comparing the baseline, new model, and fine-tuned model for epidermal growth factor receptor (EGFR). Spearman *ρ* improvements of 3.41% for the new model and 1.14% for the fine-tuned model highlight the effectiveness of the self-learning method. Results are reported separately for crystal structures (first value) and Chai1-generated structures (values in parentheses).

| **Model** | **RMSE** | **MAE** | **Pearson $r$** | **Spearman $\rho$** | **% $\uparrow \Delta$ Spearman $\rho$** |
| --- | --- | --- | --- | --- | --- |
| Baseline | 0.694<br>(0.842) | 0.572<br>(0.670) | 0.702<br>(0.593) | 0.791<br>(0.665) | 0.00% (0.00%) |
| New Model | 3.132<br>(3.407) | 3.078<br>(3.334) | 0.772<br>(0.685) | 0.818<br>(0.679) | 3.41% (2.11%) |
| Control (New Model) | 3.520<br>(3.507) | 3.455<br>(3.432) | 0.663<br>(0.580) | 0.791<br>(0.668) | 0.00% (0.45%) |
| Fine Tune | 1.033<br>(1.010) | 0.911<br>(0.858) | 0.755<br>(0.665) | 0.800<br>(0.709) | 1.14% (6.62%) |
| Control (Fine Tune) | 1.172<br>(1.103) | 1.001<br>(0.900) | 0.569<br>(0.545) | 0.773<br>(0.579) | -2.28% (-12.93%) |

<sup>a</sup> The table reports Root Mean Square Error (RMSE), Mean Absolute Error (MAE), Pearson correlation coefficient ( $r$ ), Spearman rank correlation coefficient ( $\rho$ ), and the percentage increase in Spearman $\rho$ compared to the baseline (% $\uparrow \Delta$ Spearman $\rho$ ).
<sup>b</sup> Error metrics (RMSE and MAE) are reported in units of $pK_i / pK_d$ .

## References

(1) Vos, T.; Lim, S. S.; Abbafati, C.; Abbas, K. M.; Abbasi, M.; Abbasifard, M.; Abbasi-Kangevari, M.; Abbastabar, H.; Abd-Allah, F.; Abdelalim, A.; et al. Global burden of 369 diseases and injuries in 204 countries and territories, 1990–2019: a systematic analysis for the Global Burden of Disease Study 2019. The Lancet 2020, 396 (10258), 1204–1222. DOI: 10.1016/S0140-6736(20)30925-9.

(2) Li, W.; Li, H.-L.; Wang, J.-Z.; Liu, R.; Wang, X. Abnormal protein post-translational modifications induces aggregation and abnormal deposition of protein, mediating neurodegenerative diseases. Cell & Bioscience 2024, 14 (1), 22. DOI: 10.1186/s13578-023-01189-y.

(3) Hanahan, D.; Weinberg, R. A. Hallmarks of cancer: the next generation. Cell 2011, 144 (5), 646–674. DOI: 10.1016/j.cell.2011.02.013 From NLM.

(4) Santos, R.; Ursu, O.; Gaulton, A.; Bento, A. P.; Donadi, R. S.; Bologa, C. G.; Karlsson, A.; Al-Lazikani, B.; Hersey, A.; Oprea, T. I.;, et al. A comprehensive map of molecular drug targets. Nature Reviews Drug Discovery 2017, 16 (1), 19–34. DOI: 10.1038/nrd.2016.230.

(5) Chan, J. N. Y.; Nislow, C.; Emili, A. Recent advances and method development for drug target identification. Trends in Pharmacological Sciences 2010, 31 (2), 82–88. DOI: 10.1016/j.tips.2009.11.002 (acccessed 2024/11/12).

(6) Smith, C. Drug target validation: Hitting the target. Nature 2003, 422 (6929), 342–345. DOI: 10.1038/422341a.

(7) Ashraf, S. N.; Blackwell, J. H.; Holdgate, G. A.; Lucas, S. C. C.; Solovyeva, A.; Storer, R. I.; Whitehurst, B. C. Hit me with your best shot: Integrated hit discovery for the next generation of drug targets. Drug Discovery Today 2024, 29 (10), 104143. DOI: 10.1016/j.drudis.2024.104143.

(8) Hughes, J.; Rees, S.; Kalindjian, S.; Philpott, K. Principles of early drug discovery. British Journal of Pharmacology 2011, 162 (6), 1239–1249. DOI: 10.1111/j.1476-5381.2010.01127.x.

(9) Zhang, D.; Luo, G.; Ding, X.; Lu, C. Preclinical experimental models of drug metabolism and disposition in drug discovery and development. Acta Pharmaceutica Sinica B 2012, 2 (6), 549–561. DOI: 10.1016/j.apsb.2012.10.004.

(10) Mohs, R. C.; Greig, N. H. Drug discovery and development: Role of basic biological research. Alzheimer’s & Dementia: Translational Research & Clinical Interventions 2017, 3 (4), 651–657. DOI: 10.1016/j.trci.2017.10.005.

(11) Abramson, J.; Adler, J.; Dunger, J.; Evans, R.; Green, T.; Pritzel, A.; Ronneberger, O.; Willmore, L.; Ballard, A. J.; Bambrick, J.;, et al. Accurate structure prediction of biomolecular interactions with AlphaFold 3. Nature 2024, 630 (8016), 493–500. DOI: 10.1038/s41586-024-07487-w.

(12) Boitreaud, J.; Dent, J.; McPartlon, M.; Meier, J.; Reis, V.; Rogozhonikov, A.; Wu, K. Chai-1: Decoding the molecular interactions of life. bioRxiv 2024, 2024.2010.2010.615955. DOI: 10.1101/2024.10.10.615955.

(13) Corso, G.; Deng, A.; Fry, B.; Polizzi, N.; Barzilay, R.; Jaakkola, T. Deep Confident Steps to New Pockets: Strategies for Docking Generalization. 2024; p arXiv:2402.18396.

(14) Walters, W. P.; Barzilay, R. Applications of Deep Learning in Molecule Generation and Molecular Property Prediction. Accounts of Chemical Research 2021, 54 (2), 263–270. DOI: 10.1021/acs.accounts.0c00699.

(15) Kyro, G. W.; Martin, M. T.; Watt, E. D.; Batista, V. S. CardioGenAI: A Machine Learning-Based Framework for Re-Engineering Drugs for Reduced hERG Liability. 2024; p arXiv:2403.07632.

(16) Niazi, S. K.; Mariam, Z. Computer-Aided Drug Design and Drug Discovery: A Prospective Analysis. Pharmaceuticals 2024, 17 (1), 22.

(17) Ballester, P. J.; Mitchell, J. B. A machine learning approach to predicting protein–ligand binding affinity with applications to molecular docking. Bioinformatics 2010, 26 (9), 1169–1175.

(18) Boyles, F.; Deane, C. M.; Morris, G. M. Learning from the ligand: using ligand-based features to improve binding affinity prediction. Bioinformatics 2020, 36 (3), 758–764.

(19) Zilian, D.; Sotriffer, C. A. Sfcscore rf: a random forest-based scoring function for improved affinity prediction of protein–ligand complexes. Journal of chemical information and modeling 2013, 53 (8), 1923–1933.

(20) Macari, G.; Toti, D.; Pasquadibisceglie, A.; Polticelli, F. DockingApp RF: a state-of-the-art novel scoring function for molecular docking in a user-friendly interface to AutoDock Vina. International Journal of Molecular Sciences 2020, 21 (24), 9548.

(21) Durrant, J. D.; McCammon, J. A. NNScore: a neural-network-based scoring function for the characterization of protein− ligand complexes. Journal of chemical information and modeling 2010, 50 (10), 1865–1871.

(22) Wang, H. Prediction of protein–ligand binding affinity via deep learning models. Briefings in Bioinformatics 2024, 25 (2). DOI: 10.1093/bib/bbae081 (acccessed 11/12/2024).

(23) Cang, Z.; Wei, G.-W. TopologyNet: Topology based deep convolutional and multi-task neural networks for biomolecular property predictions. PLoS computational biology 2017, 13 (7), e1005690.

(24) Öztürk, H.; Özgür, A.; Ozkirimli, E. DeepDTA: deep drug–target binding affinity prediction. Bioinformatics 2018, 34 (17), i821–i829.

(25) Zheng, L.; Fan, J.; Mu, Y. Onionnet: a multiple-layer intermolecular-contact-based convolutional neural network for protein–ligand binding affinity prediction. ACS omega 2019, 4 (14), 15956–15965.

(26) Wang, Z.; Zheng, L.; Liu, Y.; Qu, Y.; Li, Y.-Q.; Zhao, M.; Mu, Y.; Li, W. OnionNet-2: a convolutional neural network model for predicting protein-ligand binding affinity based on residue-atom contacting shells. Frontiers in chemistry 2021, 9, 753002.

(27) Wang, S.; Liu, D.; Ding, M.; Du, Z.; Zhong, Y.; Song, T.; Zhu, J.; Zhao, R. SE-OnionNet: a convolution neural network for protein–ligand binding affinity prediction. Frontiers in Genetics 2021, 11, 607824.

(28) Wang, X.; Liu, D.; Zhu, J.; Rodriguez-Paton, A.; Song, T. CSConv2d: a 2-D structural convolution neural network with a channel and spatial attention mechanism for protein-ligand binding affinity prediction. Biomolecules 2021, 11 (5), 643.

(29) Gomes, J.; Ramsundar, B.; Feinberg, E. N.; Pande, V. S. Atomic convolutional networks for predicting protein-ligand binding affinity. arXiv preprint arXiv:1703.10603 2017.

(30) Jiménez, J.; Skalic, M.; Martinez-Rosell, G.; De Fabritiis, G. K deep: protein–ligand absolute binding affinity prediction via 3d-convolutional neural networks. Journal of chemical information and modeling 2018, 58 (2), 287–296.

(31) Stepniewska-Dziubinska, M. M.; Zielenkiewicz, P.; Siedlecki, P. Development and evaluation of a deep learning model for protein–ligand binding affinity prediction. Bioinformatics 2018, 34 (21), 3666–3674.

(32) Kwon, Y.; Shin, W.-H.; Ko, J.; Lee, J. AK-score: accurate protein-ligand binding affinity prediction using an ensemble of 3D-convolutional neural networks. International journal of molecular sciences 2020, 21 (22), 8424.

(33) Kyro, G. W.; Brent, R. I.; Batista, V. S. Hac-net: A hybrid attention-based convolutional neural network for highly accurate protein–ligand binding affinity prediction. Journal of Chemical Information and Modeling 2023, 63 (7), 1947–1960.

(34) Wang, Y.; Wei, Z.; Xi, L. Sfcnn: a novel scoring function based on 3D convolutional neural network for accurate and stable protein–ligand affinity prediction. BMC bioinformatics 2022, 23 (1), 222.

(35) Jones, D.; Kim, H.; Zhang, X.; Zemla, A.; Stevenson, G.; Bennett, W. D.; Kirshner, D.; Wong, S. E.; Lightstone, F. C.; Allen, J. E. Improved protein–ligand binding affinity prediction with structure-based deep fusion inference. Journal of chemical information and modeling 2021, 61 (4), 1583–1592.

(36) Li, S.; Zhou, J.; Xu, T.; Huang, L.; Wang, F.; Xiong, H.; Huang, W.; Dou, D.; Xiong, H. Structure-aware Interactive Graph Neural Networks for the Prediction of Protein-Ligand Binding Affinity. In Proceedings of the 27th ACM SIGKDD Conference on Knowledge Discovery & Data Mining, Virtual Event, Singapore; 2021.

(37) Wang, K.; Zhou, R.; Tang, J.; Li, M. GraphscoreDTA: optimized graph neural network for protein–ligand binding affinity prediction. Bioinformatics 2023, 39 (6). DOI: 10.1093/bioinformatics/btad340 (acccessed 11/12/2024).

(38) Zhang, X.; Gao, H.; Wang, H.; Chen, Z.; Zhang, Z.; Chen, X.; Li, Y.; Qi, Y.; Wang, R. PLANET: A Multi-objective Graph Neural Network Model for Protein–Ligand Binding Affinity Prediction. Journal of Chemical Information and Modeling 2024, 64 (7), 2205–2220. DOI: 10.1021/acs.jcim.3c00253.

(39) Yang, Z.; Zhong, W.; Lv, Q.; Dong, T.; Yu-Chian Chen, C. Geometric Interaction Graph Neural Network for Predicting Protein–Ligand Binding Affinities from 3D Structures (GIGN). The Journal of Physical Chemistry Letters 2023, 14 (8), 2020–2033. DOI: 10.1021/acs.jpclett.2c03906.

(40) Li, S.; Zhou, J.; Xu, T.; Huang, L.; Wang, F.; Xiong, H.; Huang, W.; Dou, D.; Xiong, H. GIaNt: Protein-Ligand Binding Affinity Prediction via Geometry-Aware Interactive Graph Neural Network. IEEE Transactions on Knowledge and Data Engineering 2024, 36 (5), 1991–2008. DOI: 10.1109/TKDE.2023.3314502.

(41) Son, J.; Kim, D. Development of a graph convolutional neural network model for efficient prediction of protein-ligand binding affinities. PLOS ONE 2021, 16 (4), e0249404. DOI: 10.1371/journal.pone.0249404.

(42) Mastropietro, A.; Pasculli, G.; Bajorath, J. Learning characteristics of graph neural networks predicting protein–ligand affinities. Nature Machine Intelligence 2023, 5 (12), 1427–1436. DOI: 10.1038/s42256-023-00756-9.

(43) Xia, C.; Feng, S.-H.; Xia, Y.; Pan, X.; Shen, H.-B. Leveraging scaffold information to predict protein–ligand binding affinity with an empirical graph neural network. Briefings in Bioinformatics 2023, 24 (1). DOI: 10.1093/bib/bbac603 (acccessed 11/12/2024).

(44) Knutson, C.; Bontha, M.; Bilbrey, J. A.; Kumar, N. Decoding the protein–ligand interactions using parallel graph neural networks. Scientific Reports 2022, 12 (1), 7624. DOI: 10.1038/s41598-022-10418-2.

(45) Wu, J.; Chen, H.; Cheng, M.; Xiong, H. CurvAGN: Curvature-based Adaptive Graph Neural Networks for Predicting Protein-Ligand Binding Affinity. BMC Bioinformatics 2023, 24 (1), 378. DOI: 10.1186/s12859-023-05503-w.

(46) Shen, H.; Zhang, Y.; Zheng, C.; Wang, B.; Chen, P. A Cascade Graph Convolutional Network for Predicting Protein–Ligand Binding Affinity. International Journal of Molecular Sciences 2021, 22 (8), 4023.

(47) Yi, Y.; Wan, X.; Zhao, K.; Ou-Yang, L.; Zhao, P. Equivariant Line Graph Neural Network for Protein-Ligand Binding Affinity Prediction. IEEE Journal of Biomedical and Health Informatics 2024, 28 (7), 4336–4347. DOI: 10.1109/JBHI.2024.3383245.

(48) Nikolaienko, T.; Gurbych, O.; Druchok, M. Complex machine learning model needs complex testing: Examining predictability of molecular binding affinity by a graph neural network. Journal of Computational Chemistry 2022, 43 (10), 728–739. DOI: 10.1002/jcc.26831.

(49) Zhang, H.; Saravanan, K. M.; Zhang, J. Z. H. DeepBindGCN: Integrating Molecular Vector Representation with Graph Convolutional Neural Networks for Protein–Ligand Interaction Prediction. Molecules 2023, 28 (12), 4691.

(50) Yang, Y.; Zhang, R.; Lin, Z. Enhancing protein-ligand binding affinity prediction through sequential fusion of graph and convolutional neural networks. Journal of Computational Chemistry 2024, 45 (32), 2929–2940. DOI: 10.1002/jcc.27499.

(51) Yang, Z.; Zhong, W.; Lv, Q.; Dong, T.; Chen, G.; Chen, C. Y. C. Interaction-Based Inductive Bias in Graph Neural Networks: Enhancing Protein-Ligand Binding Affinity Predictions From 3D Structures. IEEE Transactions on Pattern Analysis and Machine Intelligence 2024, 46 (12), 8191–8208. DOI: 10.1109/TPAMI.2024.3400515.

(52) Mqawass, G.; Popov, P. graphLambda: Fusion Graph Neural Networks for Binding Affinity Prediction. Journal of Chemical Information and Modeling 2024, 64 (7), 2323–2330. DOI: 10.1021/acs.jcim.3c00771.

(53) Jiao, Q.; Qiu, Z.; Wang, Y.; Chen, C.; Yang, Z.; Cui, X. Edge-Gated Graph Neural Network for Predicting Protein-Ligand Binding Affinities. In 2021 IEEE International Conference on Bioinformatics and Biomedicine (BIBM), 9-12 Dec. 2021, 2021; pp 334–339. DOI: 10.1109/BIBM52615.2021.9669846.

(54) Guo, J. Improving structure-based protein-ligand affinity prediction by graph representation learning and ensemble learning. PLOS ONE 2024, 19 (1), e0296676. DOI: 10.1371/journal.pone.0296676.

(55) Nguyen, T.; Le, H.; Quinn, T. P.; Nguyen, T.; Le, T. D.; Venkatesh, S. GraphDTA: predicting drug–target binding affinity with graph neural networks. Bioinformatics 2020, 37 (8), 1140–1147. DOI: 10.1093/bioinformatics/btaa921 (acccessed 11/12/2024).

(56) Gale-Day, Z. J.; Shub, L.; Chuang, K. V.; Keiser, M. J. Proximity Graph Networks: Predicting Ligand Affinity with Message Passing Neural Networks. Journal of Chemical Information and Modeling 2024, 64 (14), 5439–5450. DOI: 10.1021/acs.jcim.4c00311.

(57) Yang, Z.; Zhong, W.; Zhao, L.; Yu-Chian Chen, C. MGraphDTA: deep multiscale graph neural network for explainable drug–target binding affinity prediction. Chemical Science 2022, 13 (3), 816–833, 10.1039/D1SC05180F. DOI: 10.1039/D1SC05180F.

(58) Jiang, M.; Wang, S.; Zhang, S.; Zhou, W.; Zhang, Y.; Li, Z. Sequence-based drug-target affinity prediction using weighted graph neural networks. BMC Genomics 2022, 23 (1), 449. DOI: 10.1186/s12864-022-08648-9.

(59) Monteiro, N. R. C.; Oliveira, J. L.; Arrais, J. P. DTITR: End-to-end drug–target binding affinity prediction with transformers. Computers in Biology and Medicine 2022, 147, 105772. DOI: 10.1016/j.compbiomed.2022.105772.

(60) Chen, D.; Liu, J.; Wei, G.-W. Multiscale topology-enabled structure-to-sequence transformer for protein–ligand interaction predictions. Nature Machine Intelligence 2024, 6 (7), 799–810. DOI: 10.1038/s42256-024-00855-1.

(61) Monteiro, N. R. C.; Oliveira, J. L.; Arrais, J. P. TAG-DTA: Binding-region-guided strategy to predict drug-target affinity using transformers. Expert Systems with Applications 2024, 238, 122334. DOI: 10.1016/j.eswa.2023.122334.

(62) Rose, T.; Monti, N.; Anand, N.; Shen, T. PLAPT: Protein-Ligand Binding Affinity Prediction Using Pretrained Transformers. bioRxiv 2024, 2024.2002.2008.575577. DOI: 10.1101/2024.02.08.575577.

(63) Shen, C.; Zhang, X.; Deng, Y.; Gao, J.; Wang, D.; Xu, L.; Pan, P.; Hou, T.; Kang, Y. Boosting Protein–Ligand Binding Pose Prediction and Virtual Screening Based on Residue–Atom Distance Likelihood Potential and Graph Transformer. Journal of Medicinal Chemistry 2022, 65 (15), 10691–10706. DOI: 10.1021/acs.jmedchem.2c00991.

(64) Zhou, C.; Li, Z.; Song, J.; Xiang, W. TransVAE-DTA: Transformer and variational autoencoder network for drug-target binding affinity prediction. Computer Methods and Programs in Biomedicine 2024, 244, 108003. DOI: 10.1016/j.cmpb.2023.108003.

(65) Han, K.; Shi, C.; Wang, Z.; Liu, W.; Li, Z.; Wang, Z.; Lei, L.; Dai, R.; Wang, M.; Zhang, Z.;, et al. Innovative Mamba and graph transformer framework for superior protein-ligand affinity prediction. Microchemical Journal 2024, 206, 111444. DOI: 10.1016/j.microc.2024.111444.

(66) Amine, A. M. E.; Fadila, A. Transformer neural network for protein-specific drug discovery and validation using QSAR. Journal of Proteins and Proteomics 2023, 14 (4), 253–262. DOI: 10.1007/s42485-023-00124-6.

(67) Vasan, A.; Gokdemir, O.; Brace, A.; Ramanathan, A.; Brettin, T.; Stevens, R.; Vishwanath, V. High Performance Binding Affinity Prediction with a Transformer-Based Surrogate Model. In 2024 IEEE International Parallel and Distributed Processing Symposium Workshops (IPDPSW), 27-31 May 2024, 2024; pp 571–580. DOI: 10.1109/IPDPSW63119.2024.00114.

(68) Tian, C.; Wang, L.; Cui, Z.; Wu, H. GTAMP-DTA: Graph transformer combined with attention mechanism for drug-target binding affinity prediction. Computational Biology and Chemistry 2024, 108, 107982. DOI: 10.1016/j.compbiolchem.2023.107982.

(69) Wang, Y.; Wu, S.; Duan, Y.; Huang, Y. A point cloud-based deep learning strategy for protein– ligand binding affinity prediction. Briefings in Bioinformatics 2021, 23 (1). DOI: 10.1093/bib/bbab474 (acccessed 11/13/2024).

(70) Wang, J.; Hu, J.; Sun, H.; Xu, M.; Yu, Y.; Liu, Y.; Cheng, L. MGPLI: exploring multigranular representations for protein–ligand interaction prediction. Bioinformatics 2022, 38 (21), 4859–4867. DOI: 10.1093/bioinformatics/btac597 (acccessed 11/13/2024).

(71) Tang, X.; Zhou, Y.; Yang, M.; Li, W. TC-DTA: Predicting Drug-Target Binding Affinity With Transformer and Convolutional Neural Networks. IEEE Transactions on NanoBioscience 2024, 23 (4), 572–578. DOI: 10.1109/TNB.2024.3441590.

(72) Li, Q.; Zhang, X.; Wu, L.; Bo, X.; He, S.; Wang, S. PLA-MoRe: A Protein–Ligand Binding Affinity Prediction Model via Comprehensive Molecular Representations. Journal of Chemical Information and Modeling 2022, 62 (18), 4380–4390. DOI: 10.1021/acs.jcim.2c00960.

(73) Xu, S.; Shen, L.; Zhang, M.; Jiang, C.; Zhang, X.; Xu, Y.; Liu, J.; Liu, X. Surface-based multimodal protein–ligand binding affinity prediction. Bioinformatics 2024, 40 (7). DOI: 10.1093/bioinformatics/btae413 (acccessed 11/13/2024).

(74) Yan, X.; Liu, Y. Graph–sequence attention and transformer for predicting drug–target affinity. RSC Advances 2022, 12 (45), 29525–29534, 10.1039/D2RA05566J. DOI: 10.1039/D2RA05566J.

(75) Wu, H.; Liu, J.; Jiang, T.; Zou, Q.; Qi, S.; Cui, Z.; Tiwari, P.; Ding, Y. AttentionMGT-DTA: A multi-modal drug-target affinity prediction using graph transformer and attention mechanism. Neural Networks 2024, 169, 623–636. DOI: 10.1016/j.neunet.2023.11.018.

(76) Liu, Y.; Xing, L.; Zhang, L.; Cai, H.; Guo, M. GEFormerDTA: drug target affinity prediction based on transformer graph for early fusion. Scientific Reports 2024, 14 (1), 7416. DOI: 10.1038/s41598-024-57879-1.

(77) Saadat, M.; Behjati, A.; Zare-Mirakabad, F.; Gharaghani, S. Drug-Target Binding Affinity Prediction Using Transformers. bioRxiv 2022, 2021.2009.2030.462610. DOI: 10.1101/2021.09.30.462610.

(78) Sun, X.; Huang, J.; Fang, Y.; Jin, Y.; Wu, J.; Wang, G.; Jia, J. MREDTA: A BERT and transformer-based molecular representation encoder for predicting drug-target binding affinity. The FASEB Journal 2024, 38 (19), e70083. DOI: 10.1096/fj.202401254R.

(79) Li, Z.; Ren, P.; Yang, H.; Zheng, J.; Bai, F. TEFDTA: a transformer encoder and fingerprint representation combined prediction method for bonded and non-bonded drug–target affinities. Bioinformatics 2023, 40 (1). DOI: 10.1093/bioinformatics/btad778 (acccessed 11/13/2024).

(80) Hu, F.; Hu, Y.; Zhang, J.; Wang, D.; Yin, P. Structure Enhanced Protein-Drug Interaction Prediction using Transformer and Graph Embedding. In 2020 IEEE International Conference on Bioinformatics and Biomedicine (BIBM), 16-19 Dec. 2020, 2020; pp 1010–1014. DOI: 10.1109/BIBM49941.2020.9313456.

(81) Quan, L.; Wu, J.; Jiang, Y.; Pan, D.; Qiang, L. DTA-GTOmega: Enhancing Drug-Target Binding Affinity Prediction with Graph Transformers Using OmegaFold Protein Structures. Journal of Molecular Biology 2024, 168843. DOI: 10.1016/j.jmb.2024.168843.

(82) Liu, S.; Wang, Y.; Deng, Y.; He, L.; Shao, B.; Yin, J.; Zheng, N.; Liu, T.-Y.; Wang, T. Improved drug–target interaction prediction with intermolecular graph transformer. Briefings in Bioinformatics 2022, 23 (5). DOI: 10.1093/bib/bbac162 (acccessed 11/13/2024).

(83) Gorantla, R.; Kubincová, A.; Suutari, B.; Cossins, B. P.; Mey, A. S. J. S. Benchmarking Active Learning Protocols for Ligand-Binding Affinity Prediction. Journal of Chemical Information and Modeling 2024, 64 (6), 1955–1965. DOI: 10.1021/acs.jcim.4c00220.

(84) Wang, R.; Fang, X.; Lu, Y.; Wang, S. The PDBbind Database: Collection of Binding Affinities for Protein−Ligand Complexes with Known Three-Dimensional Structures. Journal of Medicinal Chemistry 2004, 47 (12), 2977–2980. DOI: 10.1021/jm030580l.

(85) Wang, R.; Fang, X.; Lu, Y.; Yang, C. Y.; Wang, S. The PDBbind database: methodologies and updates. J Med Chem 2005, 48 (12), 4111–4119. DOI: 10.1021/jm048957q From NLM.

(86) Liu, Z.; Li, Y.; Han, L.; Li, J.; Liu, J.; Zhao, Z.; Nie, W.; Liu, Y.; Wang, R. PDB-wide collection of binding data: current status of the PDBbind database. Bioinformatics 2014, 31 (3), 405–412. DOI: 10.1093/bioinformatics/btu626 (acccessed 11/14/2024).

(87) Liu, Z.; Su, M.; Han, L.; Liu, J.; Yang, Q.; Li, Y.; Wang, R. Forging the Basis for Developing Protein–Ligand Interaction Scoring Functions. Accounts of Chemical Research 2017, 50 (2), 302–309. DOI: 10.1021/acs.accounts.6b00491.

(88) Su, M.; Yang, Q.; Du, Y.; Feng, G.; Liu, Z.; Li, Y.; Wang, R. Comparative Assessment of Scoring Functions: The CASF-2016 Update. Journal of Chemical Information and Modeling 2019, 59 (2), 895–913. DOI: 10.1021/acs.jcim.8b00545.

(89) Volkov, M.; Turk, J.-A.; Drizard, N.; Martin, N.; Hoffmann, B.; Gaston-Mathé, Y.; Rognan, D. On the Frustration to Predict Binding Affinities from Protein–Ligand Structures with Deep Neural Networks. Journal of Medicinal Chemistry 2022, 65 (11), 7946–7958. DOI: 10.1021/acs.jmedchem.2c00487.

(90) Li, J.; Guan, X.; Zhang, O.; Sun, K.; Wang, Y.; Bagni, D.; Head-Gordon, T. Leak Proof PDBBind: A Reorganized Dataset of Protein-Ligand Complexes for More Generalizable Binding Affinity Prediction. ArXiv 2024. From NLM.

(91) Dice, L. R. Measures of the Amount of Ecologic Association Between Species. Ecology 1945, 26 (3), 297–302. DOI: 10.2307/1932409 (acccessed 2024/11/14/).JSTOR.

(92) Needleman, S. B.; Wunsch, C. D. A general method applicable to the search for similarities in the amino acid sequence of two proteins. J Mol Biol 1970, 48 (3), 443–453. DOI: 10.1016/0022-2836(70)90057-4 From NLM.

(93) Wang, D. D.; Xie, H.; Yan, H. Proteo-chemometrics interaction fingerprints of protein-ligand complexes predict binding affinity. Bioinformatics 2021, 37 (17), 2570–2579. DOI: 10.1093/bioinformatics/btab132 From NLM.

(94) Bajusz, D.; Rácz, A.; Héberger, K. Why is Tanimoto index an appropriate choice for fingerprint-based similarity calculations? Journal of Cheminformatics 2015, 7 (1), 20. DOI: 10.1186/s13321-015-0069-3.

(95) Liu, T.; Lin, Y.; Wen, X.; Jorissen, R. N.; Gilson, M. K. BindingDB: a web-accessible database of experimentally determined protein–ligand binding affinities. Nucleic Acids Research 2006, 35 (suppl_1), D198–D201. DOI: 10.1093/nar/gkl999 (acccessed 11/14/2024).

(96) Burley, S. K.; Bhikadiya, C.; Bi, C.; Bittrich, S.; Chao, H.; Chen, L.; Craig, P. A.; Crichlow, G. V.; Dalenberg, K.; Duarte, J. M.;, et al. RCSB Protein Data Bank (RCSB.org): delivery of experimentally-determined PDB structures alongside one million computed structure models of proteins from artificial intelligence/machine learning. Nucleic Acids Research 2022, 51 (D1), D488–D508. DOI: 10.1093/nar/gkac1077 (acccessed 11/14/2024).

(97) Gao, K.; Wang, R.; Chen, J.; Tepe, J. J.; Huang, F.; Wei, G.-W. Perspectives on SARS-CoV-2 Main Protease Inhibitors. Journal of Medicinal Chemistry 2021, 64 (23), 16922–16955. DOI: 10.1021/acs.jmedchem.1c00409.

(98) Herbst, R. S. Review of epidermal growth factor receptor biology. Int J Radiat Oncol Biol Phys 2004, 59 (2 Suppl), 21–26. DOI: 10.1016/j.ijrobp.2003.11.041 From NLM.

(99) Kyro, G. W.; Morgunov, A.; Brent, R. I.; Batista, V. S. ChemSpaceAL: An Efficient Active Learning Methodology Applied to Protein-Specific Molecular Generation. Journal of Chemical Information and Modeling 2024, 64 (3), 653–665. DOI: 10.1021/acs.jcim.3c01456.

(100) Mendez, D.; Gaulton, A.; Bento, A. P.; Chambers, J.; De Veij, M.; Félix, E.; Magariños, M. P.; Mosquera, J. F.; Mutowo, P.; Nowotka, M. ChEMBL: towards direct deposition of bioassay data. Nucleic acids research 2019, 47 (D1), D930–D940.

(101) Brown, N.; Fiscato, M.; Segler, M. H.; Vaucher, A. C. GuacaMol: benchmarking models for de novo molecular design. Journal of chemical information and modeling 2019, 59 (3), 1096–1108.

(102) Polykovskiy, D.; Zhebrak, A.; Sanchez-Lengeling, B.; Golovanov, S.; Tatanov, O.; Belyaev, S.; Kurbanov, R.; Artamonov, A.; Aladinskiy, V.; Veselov, M. Molecular sets (MOSES): a benchmarking platform for molecular generation models. Frontiers in pharmacology 2020, 11, 565644.

(103) Cock, P. J.; Antao, T.; Chang, J. T.; Chapman, B. A.; Cox, C. J.; Dalke, A.; Friedberg, I.; Hamelryck, T.; Kauff, F.; Wilczynski, B.;, et al. Biopython: freely available Python tools for computational molecular biology and bioinformatics. Bioinformatics 2009, 25 (11), 1422–1423. DOI: 10.1093/bioinformatics/btp163 From NLM.

(104) Eastman, P.; Swails, J.; Chodera, J. D.; McGibbon, R. T.; Zhao, Y.; Beauchamp, K. A.; Wang, L.-P.; Simmonett, A. C.; Harrigan, M. P.; Stern, C. D.;, et al. OpenMM 7: Rapid development of high performance algorithms for molecular dynamics. PLOS Computational Biology 2017, 13 (7), e1005659. DOI: 10.1371/journal.pcbi.1005659.

(105) O’Boyle, N. M.; Banck, M.; James, C. A.; Morley, C.; Vandermeersch, T.; Hutchison, G. R. Open Babel: An open chemical toolbox. Journal of Cheminformatics 2011, 3 (1), 33. DOI: 10.1186/1758-2946-3-33.

(106) RDKit: Open-Source Cheminformatics Software. https://www.rdkit.org (accessed 2024.

(107) Los Alamos National Laboratory, Periodic Table of Elements. https://periodic.lanl.gov/index.shtml (accessed 2024.

(108) Schwerdtfeger, P.; Nagle, J. K. 2018 Table of static dipole polarizabilities of the neutral elements in the periodic table*. Molecular Physics 2019, 117 (9-12), 1200–1225. DOI: 10.1080/00268976.2018.1535143.

(109) Sverrisson, F.; Feydy, J.; Correia, B. E.; Bronstein, M. M. Fast end-to-end learning on protein surfaces. bioRxiv 2020, 2020.2012.2028.424589. DOI: 10.1101/2020.12.28.424589.

(110) Lin, F.-Y.; Liu, C.-I.; Liu, Y.-L.; Zhang, Y.; Wang, K.; Jeng, W.-Y.; Ko, T.-P.; Cao, R.; Wang, A. H.-J.; Oldfield, E. Mechanism of action and inhibition of dehydrosqualene synthase. Proceedings of the National Academy of Sciences 2010, 107 (50), 21337–21342. DOI: doi:10.1073/pnas.1010907107.

(111) Lin, Z.; Akin, H.; Rao, R.; Hie, B.; Zhu, Z.; Lu, W.; Smetanin, N.; Verkuil, R.; Kabeli, O.; Shmueli, Y.;, et al. Evolutionary-scale prediction of atomic-level protein structure with a language model. Science 2023, 379 (6637), 1123–1130. DOI: doi:10.1126/science.ade2574.

(112) Bouysset, C.; Fiorucci, S. ProLIF: a library to encode molecular interactions as fingerprints. Journal of Cheminformatics 2021, 13 (1), 72. DOI: 10.1186/s13321-021-00548-6.

(113) Satorras, V. c. G.; Hoogeboom, E.; Welling, M. E(n) Equivariant Graph Neural Networks. In Proceedings of the 38th International Conference on Machine Learning, Proceedings of Machine Learning Research; 2021.

(114) Loshchilov, I.; Hutter, F. Decoupled Weight Decay Regularization. 2017; p arXiv:1711.05101.

(115) PyTorch Lightning; 2019. https://www.pytorchlightning.ai (accessed.

(116) Gal, Y.; Ghahramani, Z. Dropout as a Bayesian Approximation: Representing Model Uncertainty in Deep Learning. 2015; p arXiv:1506.02142.

(117) Bengio, Y.; Courville, A.; Vincent, P. Representation Learning: A Review and New Perspectives. IEEE Transactions on Pattern Analysis and Machine Intelligence 2013, 35 (8), 1798–1828. DOI: 10.1109/TPAMI.2013.50.

(118) Ben-David, S.; Blitzer, J.; Crammer, K.; Pereira, F. Analysis of Representations for Domain Adaptation. In Advances in Neural Information Processing Systems 19: Proceedings of the 2006 Conference, Schölkopf, B., Platt, J., Hofmann, T. Eds.; The MIT Press, 2007; p 0.

(119) Cang, Z.; Mu, L.; Wei, G.-W. Representability of algebraic topology for biomolecules in machine learning based scoring and virtual screening. PLOS Computational Biology 2018, 14 (1), e1005929. DOI: 10.1371/journal.pcbi.1005929.

(120) Meli, R.; Anighoro, A.; Bodkin, M. J.; Morris, G. M.; Biggin, P. C. Learning protein-ligand binding affinity with atomic environment vectors. Journal of Cheminformatics 2021, 13 (1), 59. DOI: 10.1186/s13321-021-00536-w.

(121) Li, Y.; Rezaei, M. A.; Li, C.; Li, X. DeepAtom: A Framework for Protein-Ligand Binding Affinity Prediction. In 2019 IEEE International Conference on Bioinformatics and Biomedicine (BIBM), 18-21 Nov. 2019, 2019; pp 303–310. DOI: 10.1109/BIBM47256.2019.8982964.

(122) Meng, Z.; Xia, K. Persistent spectral–based machine learning (PerSpect ML) for protein-ligand binding affinity prediction. Science Advances 2021, 7 (19), eabc5329. DOI: doi:10.1126/sciadv.abc5329.

(123) Nguyen, D. D.; Wei, G.-W. AGL-Score: Algebraic Graph Learning Score for Protein–Ligand Binding Scoring, Ranking, Docking, and Screening. Journal of Chemical Information and Modeling 2019, 59 (7), 3291–3304. DOI: 10.1021/acs.jcim.9b00334.

(124) Liu, X.; Feng, H.; Wu, J.; Xia, K. Persistent spectral hypergraph based machine learning (PSH-ML) for protein-ligand binding affinity prediction. Briefings in Bioinformatics 2021, 22 (5), bbab127.

(125) Seo, S.; Choi, J.; Park, S.; Ahn, J. Binding affinity prediction for protein–ligand complex using deep attention mechanism based on intermolecular interactions. BMC bioinformatics 2021, 22, 1–15.

(126) Wang, K.; Zhou, R.; Li, Y.; Li, M. DeepDTAF: a deep learning method to predict protein– ligand binding affinity. Briefings in Bioinformatics 2021, 22 (5), bbab072.

(127) Wang, H.; Liu, H.; Ning, S.; Zeng, C.; Zhao, Y. DLSSAffinity: protein–ligand binding affinity prediction via a deep learning model. Physical Chemistry Chemical Physics 2022, 24 (17), 10124–10133.

(128) Zhu, F.; Zhang, X.; Allen, J. E.; Jones, D.; Lightstone, F. C. Binding affinity prediction by pairwise function based on neural network. Journal of chemical information and modeling 2020, 60 (6), 2766–2772.

(129) Li, X.-S.; Liu, X.; Lu, L.; Hua, X.-S.; Chi, Y.; Xia, K. Multiphysical graph neural network (MP-GNN) for COVID-19 drug design. Briefings in Bioinformatics 2022, 23 (4). DOI: 10.1093/bib/bbac231 (acccessed 11/18/2024).

(130) Wójcikowski, M.; Kukiełka, M.; Stepniewska-Dziubinska, M. M.; Siedlecki, P. Development of a protein-ligand extended connectivity (PLEC) fingerprint and its application for binding affinity predictions. Bioinformatics 2019, 35 (8), 1334–1341. DOI: 10.1093/bioinformatics/bty757 From NLM.

(131) Paszke, A.; Gross, S.; Massa, F.; Lerer, A.; Bradbury, J.; Chanan, G.; Killeen, T.; Lin, Z.; Gimelshein, N.; Antiga, L.; et al. PyTorch: An Imperative Style, High-Performance Deep Learning Library. 2019; p arXiv:1912.01703.

(132) Fey, M.; Lenssen, J. E. Fast Graph Representation Learning with PyTorch Geometric. 2019; p arXiv:1903.02428.

(133) Charlier, B.; Feydy, J.; Glaunès, J. A.; Collin, F.-D.; Durif, G. Kernel Operations on the GPU, with Autodiff, without Memory Overflows. 2020; p arXiv:2004.11127.

(134) Virtanen, P.; Gommers, R.; Oliphant, T. E.; Haberland, M.; Reddy, T.; Cournapeau, D.; Burovski, E.; Peterson, P.; Weckesser, W.; Bright, J.;, et al. SciPy 1.0: fundamental algorithms for scientific computing in Python. Nature Methods 2020, 17 (3), 261–272. DOI: 10.1038/s41592-019-0686-2.

